# Glacier Retreat Effects On Ecosystem Development And Carbon Dynamics

**DOI:** 10.1101/2024.10.06.616884

**Authors:** Ceresa Erica, Velasquez Laura, Charles Cécile, Grand Stéphanie, Losapio Gianalberto

## Abstract

One of the clearest indicators of global warming is the retreat of Alpine glaciers. Organisms rapidly colonize bare ground free from ice, and in less than a century complex ecosystems can develop in a process of primary succession, where community changes as terrain age increases and interactions within species change over time. When glaciers retreat the resulting ice-free soil is exposed to new forms of chemical and biological weathering, with initiating soil formation as an important driver of evolving ecosystems. Soils play an important role in the global carbon (C) cycle, serving as the largest reservoir of carbon. CO_2_ is removed from the atmosphere by plant photosynthesis, then stored in biomass and soils, and released again to the atmosphere via plant and soil respiration. Soil respiration includes CO_2_ production by heterotrophic soil organisms (e.g., bacteria, fungi, invertebrates) that metabolize plant litter and soil organic matter, and autotrophs (e.g. plant roots). However, little is known about the development of soils, their physico-chemical changes, and their influence on plant diversity and soil carbon emission (CO_2_) after glacier retreat.

This study aims to investigate how the retreat of the Mont Miné and Ferpècle glaciers (Valais Alps, Switzerland) influences soil functioning and vegetation development. For this purpose, we carried out soil analyses and vegetation surveys on 5 ice-free stages deglaciated between 1864 and 2023. Soil physico-chemical properties (pH, soil organic matter, organic carbon and nitrogen, granulometry, available elements, total elements), soil CO_2_ respiration fluxes, and plant communities (composition, diversity, richness, biomass) were studied in 20 plots along the glacier forefield.

Our results show that vegetation develops according to the principles of primary succession, going further from glacier front: from pioneer vegetation 7 years after glacier retreat, progressing through early vegetation at 20 years, then to intermediate vegetation, and ultimately culminating in a climax coniferous forest, 140 years after glacier retreat. Over 140 years, the increase in vegetation productivity results in a greater transfer of organic matter (SOM) to the soil, which increases organic carbon, nitrogen, and nutrient levels, and decreases pH values as it releases organic acids. Aboveground plant biomass (i.e. carbon storage) increases over the succession, however, plant diversity and richness peak at the intermediate stages and then level off due to the dominance of few species in the late stages. At the intermediate stage, there is also a peak observed in soil respiration (CO_2_ flux), which in turn promotes plant colonization and diversity.

Thus, the soil, along with its development stages and properties, plays a critical role in carbon accumulation, leading to an increased potential for carbon release as organic matter storage increases.

This project aims to contribute to the understanding of the consequences of global warming on recently deglaciated ecosystems, which remain still poorly overlooked.

## 1. Introduction

Earth’s temperature is increasing at an unprecedented rate driven by enhanced anthropogenic greenhouse gas emissions which severely impact ecosystems. Mountain surface air temperatures in the European Alps show warming at an average of 0.3°C per decade (Hock et al., 2019). The expected future climate scenarios for Switzerland include not only warming temperatures but also drier summers, increased heavy precipitation, and snow-scarce winters (NCCS, 2018), impacting the water cycle and the glacier ice mass. Due to global warming, the loss of ice mass is occurring at an alarming pace, and the expected warmer winter temperatures will not compensate for the loss of ice. It is projected that alpine glaciers will lose at least a third of their volume by 2050, even if greenhouse gas emissions cease entirely (Cook et al., 2023). Even under the Paris 1.5°C accord, by 2100, half of the glaciers worldwide are expected to disappear (Rounce et al., 2023).

### 1.1 Consequences of glacier retreat on biodiversity and soil

The retreat of glaciers affects the biological community (Stibal et al., 2020). Although glacial environments are generally considered to be hostile, with harsh conditions, and characterized by high levels of disturbance, low nutrient availability, and low productivity, glacier melting contribution to terrestrial ecosystems is reported to create a mosaic of environmental conditions that are crucial for a large number of plants and animals (Cauvy-Fraunié & Dangles, 2019, Jacobsen & Dangles, 2012).

Indeed, during glacial retreat, the ice-free ground is exposed to colonization by organisms, where species composition changes over time in a gradient called primary succession. Within a few years, microorganisms, plants, and animals create diverse and complex communities, whose age increases with increasing distance from the retreating glacier. Four stages of primary succession occur once the glacier retreats: pioneer, early, intermediate, and late successional (Eichel et al., 2013). The trend of succession is outlined by increasing vegetation cover, biomass, and vegetation stratification with increasing age from the glacier front.

In this process of primary succession, vegetation establishment is tightly bound to the characteristics of the underlying substrate, especially in the pioneer and the early stage, where small-scale abiotic factors such as microtopography, grain size and water content of the substrate seem to be crucial for the development of both vegetation and soil (Burga et al., 2010), and as primary succession progresses, biotic processes such as competition, facilitation, tolerance, and inhibition become more significant.

Succession is a complex and dynamic process, where species interactions drive ecosystem development. Early colonizers play a crucial role by modifying the environment to improve the suitability for later succession species, a concept known as "facilitation" model (Ficetola et al., 2021). As the ecosystem progresses into intermediate and late successional stages, these early colonizers are gradually replaced by more competitive, slow-growing, and tolerant species. This transition marks a shift towards the "competition" model, where species traits such as limited dispersal and slow growth rates, which may have limited success in early colonization stages, become advantageous. Additionally, species in later successional stages often exhibit traits that might limit their success at early colonization stages (e.g. limited disperals or slow growth rate), and/or have better tolerance. Hence, identifying the biodiversity patterns along a chronosequence of glacier retreat is not straightforward, as organisms exhibit diverse responses to changes in environmental conditions (Stibal et al., 2020). Specialist species uniquely adapted to glacial conditions may disappear, losing their ecological niche and being replaced by species with great capacity for adaptation, generalist species colonizing from downstream Cauvy-Fraunié & Dangles, 2019). This shift is expected to boost the abundance and richness of generalist species, establishing themselves as dominant winners. Because of these shifts in associations plant diversity initially increases with glacier retreat, but ultimately decreases after glacier extinction (Losapio et al., 2021).

Furthermore, the loss of glaciers implies a change in the composition and development of soil as the ice-free ground is exposed to new forms of chemical and biological weathering with initiating soil formation as an important driver of evolving ecosystems.

Generally, soil development occurs through a dynamic process involving the additions and transformations of materials within the system, coupled with the effect of time. As materials such as rock fragments, minerals, organic matter, and nutrients are introduced into the soil through various processes like weathering, erosion, and biological activity, they contribute to the formation of distinct soil horizons or layers (Chapin et al., 2002). These horizons, known as O (organic), A (topsoil), E (eluviation), B (subsoil), and C (parent material), develop over time through the accumulation, leaching, and transformation of soil constituents (Chapin et al., 2002).

Soil development is tied to primary succession, serving as the substrate upon which biological communities establish. Microbes, vegetation, and fauna collectively contribute to soil development (Schulz et al., 2013) and to the accumulation of soil organic matter (SOM) (Egli et al., 2006), thus crucial for carbon storage and cycles (Egli et al., 2012.) Soil depth increases as humus from decaying plant matter and soil microbes accumulate. Soil deepening is further enhanced through the intermixing of organic material with weathered minerals, leading to the formation of organo-mineral associations that play a significant role in stabilizing soil organic matter (Schweizer et al., 2018). With the progression of plant cover and soil development, distinct soil horizons gradually form due to the ongoing processes of mineral weathering and leaching (Mavris et al., 2010). Understanding the biogeochemical properties of the soil is pivotal when considering how vegetation communities are formed, as these properties can influence the development and composition of plant communities. For instance, in proglacial areas the high rates of mineral formation, chemical and biological weathering on young surfaces lead to rapid soil development (Egli et al., 2006). In soil development and vegetation establishment the grain size of the substrate, with other soil properties, plays a crucial role, especially in the pioneer and early stages (Burga et al., 2010). As soil is weathered, the conversion of primary to secondary minerals increases the proportion of small soil particles (Chapin et al., 2002). Hence, from coarse-rich soil, at the glacier front, the soil texture becomes progressively finer (Burga et al., 2010). Through weathering processes, elements like potassium, calcium, and magnesium are slowly released from minerals, providing a source of nutrients for plants and promoting vegetation development.

As vegetation cover and productivity increase, so does the transfer of organic matter to the soil, a critical component of soils, composed of plant, animal, and microbial residues. Soil organic matter provides the energy and carbon for heterotrophic soil organisms and is an important reservoir of essential nutrients required for plant growth (Chapin et al., 2002). SOM strongly affects the rates of weathering and soil development, soil water-holding capacity, and soil structure. Moreover, plant roots play a significant role in nutrient cycling by extracting nutrients from the soil and releasing them back into the soil (Chapin et al., 2002). Microorganisms, such as bacteria and fungi, further break down organic matter, releasing nutrients in forms that plants can absorb. As vegetation cover increases, the cycling of nutrients becomes more efficient, leading to an overall rise in soil nutrient levels.

Additionally, the increase in soil carbon content promotes soil respiration. Soil respiration, also known as microbial respiration, is the process by which soil microorganisms break down organic matter and release carbon dioxide into the atmosphere. As soil organic matter and organic carbon levels rise, there is a greater availability for microbial decomposition. This leads to an increase in microbial activity, which in turn results in higher rates of soil respiration. The release of CO_2_ through soil respiration is a natural part of the carbon cycle, where carbon is cycled between the soil, plants, and the atmosphere. Another important component of the carbon cycle is the carbon stock in above-ground biomass (AGB). Quantifying the estimate of AGB along glacier retreat provides important insights into ecosystem recovery and carbon dynamics, understanding the role of vegetation succession in carbon sequestration, as well as the potential contribution of the ecosystems to mitigating climate change.

Finally, the balance between input, largely from plant-derived C source, and output, mainly through soil respiration, of carbon in soils is of great importance. If carbon losses from soils keep increasing, it may have positive feedback on climate warming by increasing CO_2_ concentration in the atmosphere (Smith et al., 2008) and rising ultimately atmospheric temperatures.

To sum up, soil development of alpine areas is a dynamic and complex process influenced by a multitude of factors, including topography, climate, vegetation, and geological history. Yet, there is still a gap in current knowledge about the linkage between glacier retreat and its influence on soil respiration processes and plant-soil interactions.

The purpose of this thesis work is to understand the intricate linkages of these factors in shaping soil properties and vegetation development of glacier forefields. By examining key soil features such as organic matter, organic carbon content, nutrient availability and element composition, pH, and soil texture, insights will be gained into the long-term processes that influence the development of soils.

### 1.2 Research objectives

Despite growing interest and research in biodiversity response to glacier retreat, the underlying processes are still uncertain, especially the interactions among vegetation, soil, and carbon flux, which remains overlooked.

The overall aim of the present study is to improve the understanding of initial soil carbon cycling along glacier forefields, combining gas exchange with soil and vegetation measurements. The objectives of the current research are the following:

- To characterize vegetation development, with plant diversity and richness, and aboveground biomass;
- To assess soil properties for quantifying soil development;
- To quantify soil carbon emissions (i.e., soil respiration) and its change with soil formation and glacier retreat;
- Finally, to assess the interaction between vegetation and soil development and the observed carbon fluxes.

Understanding the linkage between the evolution of soil, their physical and chemical changes, and their influence on plant growth in recently ice-free areas will help to gain insight into how alpine ecosystems will answer to current climate change. This understanding will also aid in modeling vegetation development following deglaciation, thereby constructing scenarios for future alpine greening. To answer these questions, studies focusing on the consequences of glacier retreat on biodiversity and potential feedback to climate change are necessary.

To address the research gaps on the development of ecosystem and carbon dynamics during initial ecosystem formation in an alpine glacier forefield, we hypothesized that, going further away from the glacier front:

- Vegetation diversity and biomass will increase, along the primary succession;
- Soil will be more developed, will be enriched in organic matter, in nutrients and carbon, granulometry will shifts towards clay-sized particles;
- Soil respiration will increase along the glacier forefield, given by the high rates of microbial activity associated with higher vegetation cover and productivity.

## 2. Materials and Methods

### 2.1 Study area

Located in the Swiss Alps, the investigated area is located in the southern part of the canton Valais, at the end of the the Hérens Valley (Figure 2). The study site is the forefield where the glacier of Mont Miné and the glacier of Ferpècle were united until the early 1960 (Nicolussi et al., 2022). Since then, both glaciers have been undergoing an unprecedented retreat, also as demonstrated by GLAMOS (Glacier Monitoring Switzerland (GLAMOS), 2023). During the period 1956-2020 Mont Miné glacier has lost 854 m of its length and Ferpècle glacier has lost 2364 m of its length since 1891 (referred time 1891-2023).

**Figure 1:**
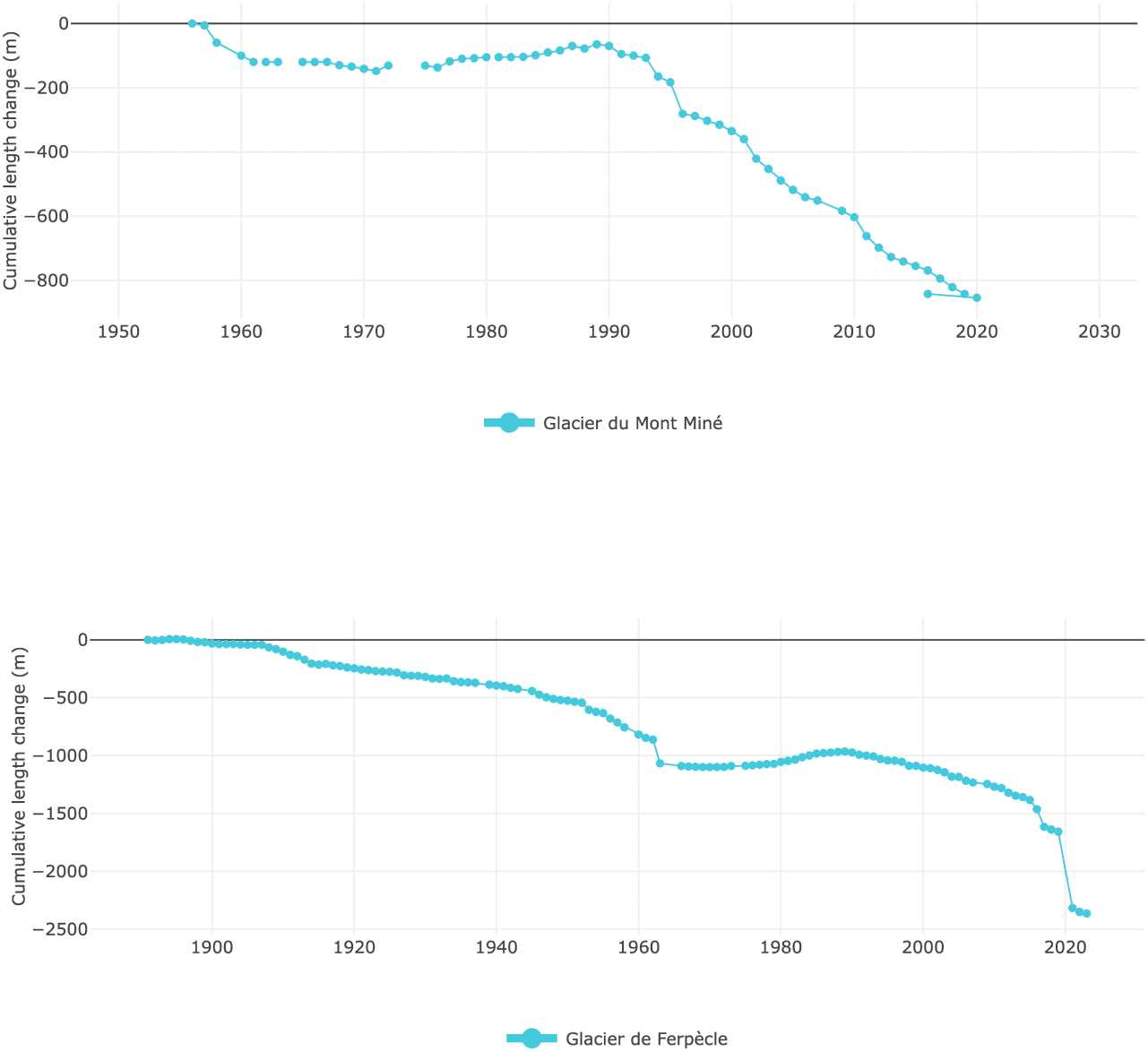
Cumulative length variation of Mont Miné Glacier and Ferpècle Glacier (*«Glaciers Suisses | Réseau des relevés glaciologiques suisse GLAMOS», 2022*)

**Figure 2:**
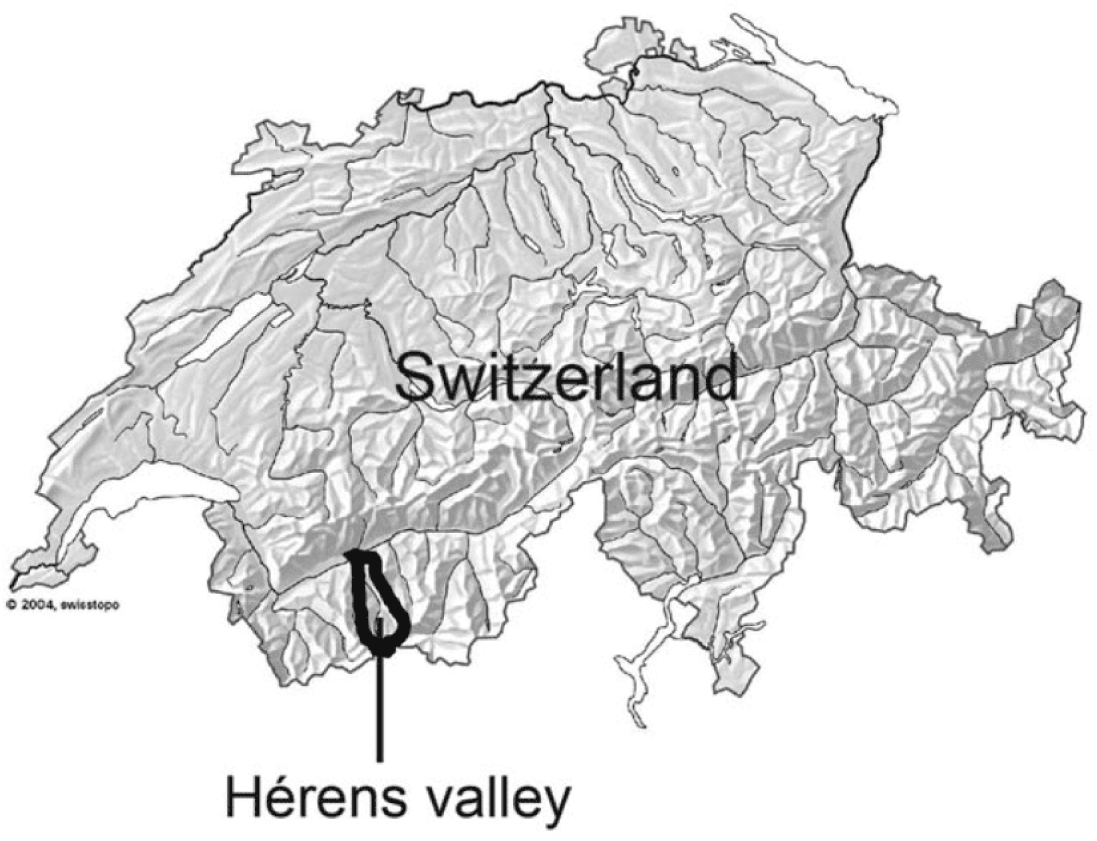
Location of the Val d’Hérens in Switzerland

The geology of the area is mainly composed of crystalline rocks such as schistous quartzites, micaschists, gneisses and amphibolites (Lambiel et al., 2016). The Valley is sheltered from the main atmospheric disturbances due to its position in the central part of the Alps. As a consequence, the climate of the valley is relatively dry, with a mean annual precipitation of 720 mm (mean 1987–2012, at 1825 m a.s.l.) and the 0°C isotherm is around 2600 m a.s.l. (Lambiel et al., 2016).

The lower altitude of the site is 1883 m a.s.l. and the highest is 2080 m a.s.l.. The low altitudinal gradient minimizes potential biases in colonization patterns resulting from significant differences in elevation as the altitudinal gradient along glacier foreland is an abiotic factor that can influence the pattern of distribution of species (Ficetola et al., 2021). The ice-free areas provide an ideal experimental model for understanding changes in ecological networks, colonization patterns, and community formation.

The area was divided in 5 stages, which were determined based on the former glacier positions and based on the position of frontal and lateral moraine. The stages correspond to the definition of stage of ecological succession of the vegetation according to Eichel et al. (Eichel, 2019). The order of the stage subdivision ranges from 0 to 4, from 0 the most recently deglaciated to stage 4, the terrains furthest downstream from the current glacier front (3).

In the study site, the moraines delimited the following stages (4):

Data regarding vegetation, soil, and soil respiration were collected in a month of fieldwork, from June the 15th to July the 15th 2023. The summer was characterized by low precipitation and high temperature (MeteoSvizzera, 2023).

After fieldwork, the soil samples were processed and analyzed in the laboratory facilities of the University of Lausanne from August 2023 to December 2023. Laboratory analysis was followed by data processing and statistical analysis.

### 2.2 Field work and sampling plan

For each stage determined in the field, 4 randomly distributed plots of 5 x 5 meters were samples to characterize vegetation diversity and biomass data, soil properties, and soil respiration.

### Vegetation

Vegetation surveys were initially conducted to gather data on diversity indices, followed by the collection of aboveground biomass (AGBest) data to estimate the carbon stock within the vegetation.

In each plot, vegetation surveys were carried out. Identification of plants was done with the help of the Flora Helvetica (Lauber et al., 2001) by a botanist. Vegetation diversity data were then used to quantify various indices and determine different ecological properties for each plot:

- **Shannon diversity index (H).** The Shannon index represents a way to measure the diversity of species in a community. Denoted as H, this index is calculated as:

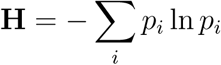

where *p* is the proportion of individuals for each species within the total number of individuals in the community.

- **Species richness.** It represents the total count of species within each plot.
- **Plant functional group.** Represents the proportions of trees, shrubs, forbs, and graminoids in each plot. This index was computed by determining the percentage of each group, which was calculated based on the weighted abundance of each species.

Regarding the biomass of trees (i.e. carbon stock in aboveground vegetation), it has been estimated using the Nikon Forestry Pro II Laser Range Finder (‘Nikon Forestry Pro II Laser Range Finder’, n.d.). This device provides simple and fast determination of relevant sizes and tree heights with its integrated clinometer.

To estimate tree biomass, we first measured the height of trees using the Nikon Forestry Pro instrument, then determined the Diameter at Breast Height (DBH) with a measuring tape, allowing us to approximate each tree to a cylinder volume. For shrubs, we measured their length and average height within the plot. Graminaceous and forb aboveground biomass was assessed by cutting samples from a 1x1 smaller plot within each plot. Once cut, they were put into bags and then left to dry in an oven. Once dried, they were weighed. Afterward, the total aboveground biomass was calculated by applying the biomass estimation formulas specific to each type (Conti et al., 2019), taking into account the density of each tree.

The equation for estimating the aboveground biomass (AGBest*_t_*) of trees is:

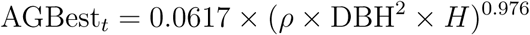

Whereas for shrubs the estimated aboveground biomass (AGBest*_s_*) is:

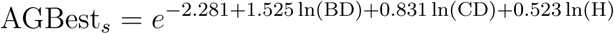

Where: AGBest = estimated aboveground biomass (kg), DBH = diameter at 130 cm stem height (cm); BD = stem basal diameter (cm); CD = mean crown diameter (m); H = height (m); *ρ* = wood density (g/cm^3^)

Estimating biomass (with the quantification of soil respiration) along a primary succession provides valuable comprehension into ecosystem dynamics and carbon cycling following glacier retreat and ecosystem development. Trees and vegetation store significant amounts of carbon in biomass, and such biomass allows estimating the amount of carbon sequestered aboveground by an ecosystem.

### Soil development

In each plot, soil samples were collected in each corner of the plot and in the center for the the first 10 cm of ground. By examining key soil properties such as organic carbon content, nutrient availability, pH, and soil texture, insights will be gained into the long-term development of soils in glacial areas.

### Soil respiration

Soil respiration was measured with the LICOR device, an automated closed transparent chamber attached to the analyzer LI 8100A (Figure 8) (Li-Cor Biosciences, Lincoln, NE, USA). The LI-800A is a fully automated system designed for measuring soil CO_2_ fluxes. The system includes an Analyzer Control Unit that powers a survey chamber, containing the system’s electronics, and an infrared gas analyzer (IRGA) for measuring changes in CO_2_ and H_2_O concentrations within the soil chamber. The LI-8100A utilizes the rate of CO_2_ increase in a measurement chamber to estimate how quickly CO_2_ diffuses into the surrounding air outside the chamber. To ensure accurate estimation, similar concentration gradients, pressure, temperature, and soil moisture must exist inside and outside the chamber. The process of calculating soil CO_2_ efflux involves fitting a linear function to the "linear portion" of the curve.

**Figure 3:**
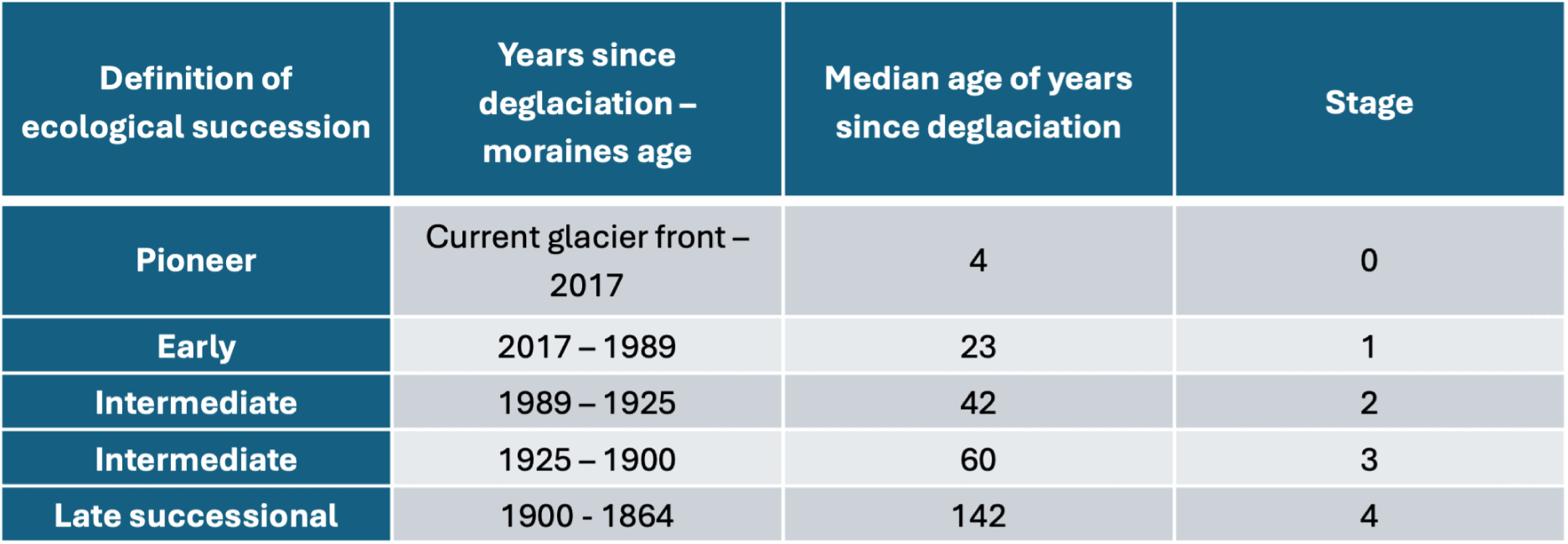
Subdivision of our study area according to the definition of ecological succession using moraine’s age as reference.

**Figure 4:**
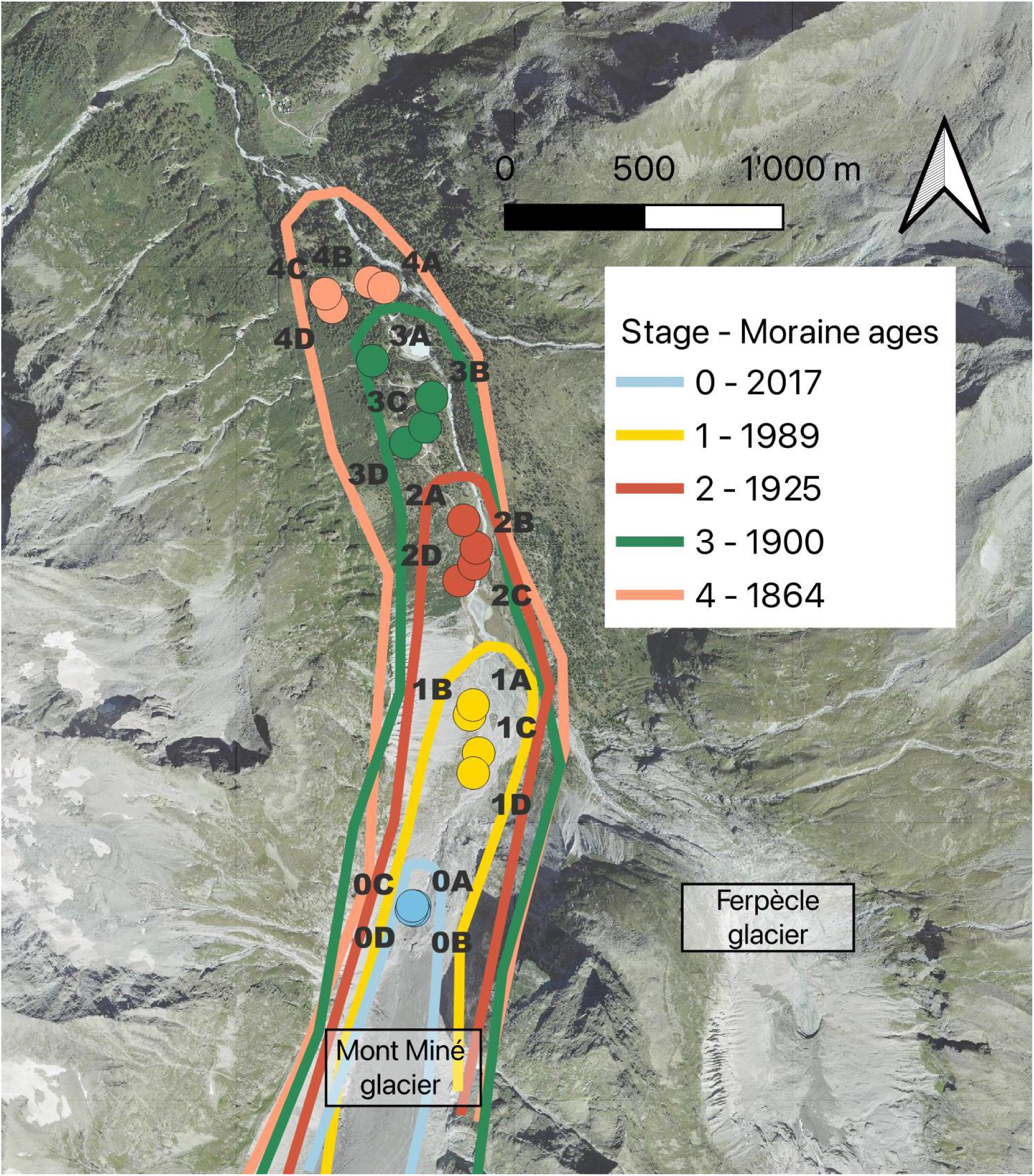
Satellite image of Vallon de Ferpècle with geochronological reconstruction of Mont Miné glacier’s moraines and respective stages of primary succession.

**Figure 5:**
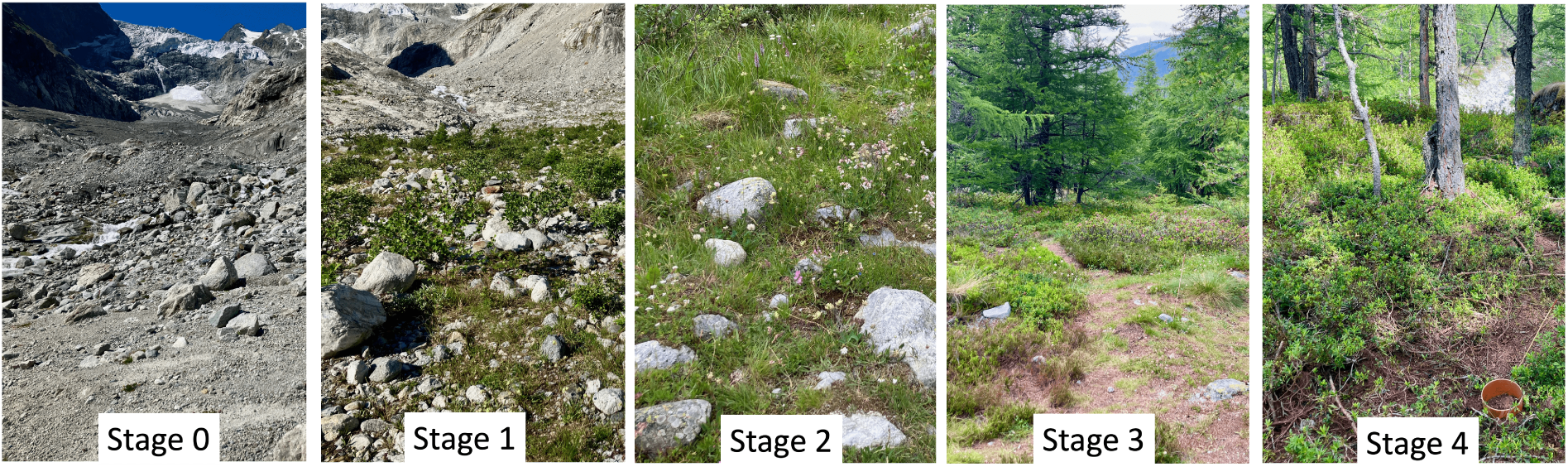
A picture for each stage of the succession to show macroscopic habitat differences.

**Figure 6:**
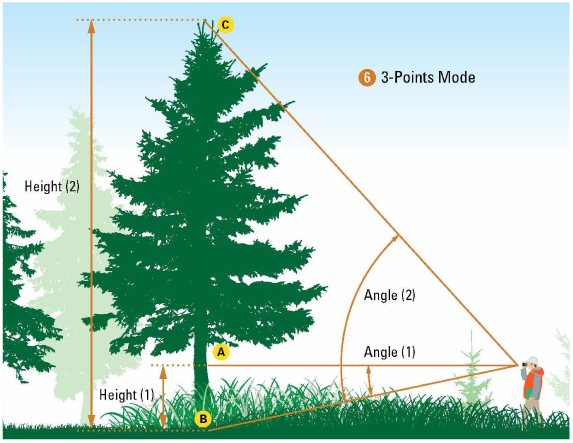
The operation of measuring tree height using the three-points method

**Figure 7:**
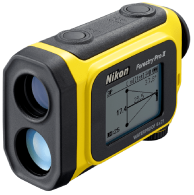
The Nikon Forestry Pro II Laser Range Finder

**Figure 8:**
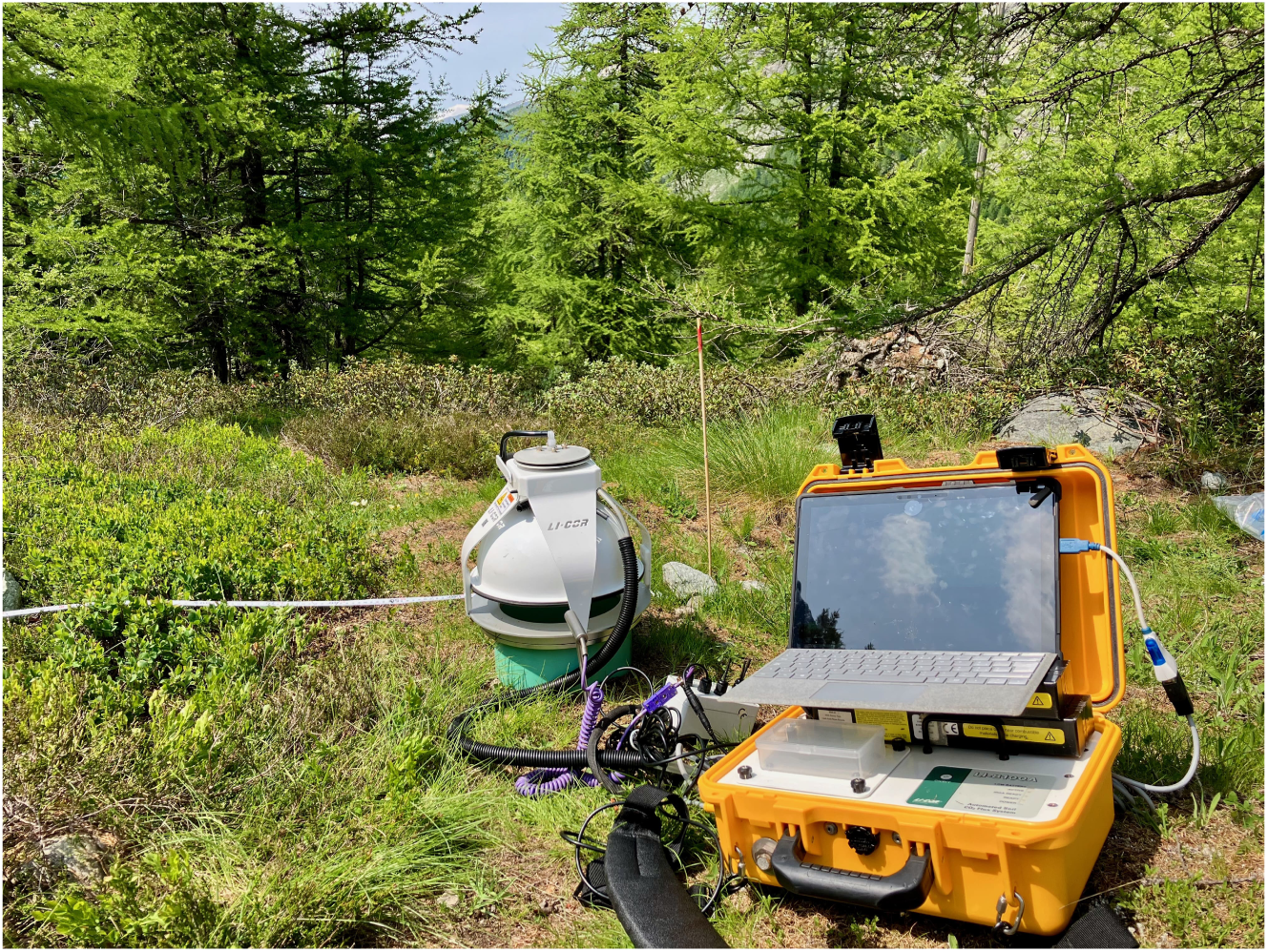
The LI-800A gas chamber used to detect soil respiration attached to the analyzer. Measurements are shown in real time on the computer through the program Soil-FluxPro™.

In summary, CO_2_ concentration is measured by the analyzer’s optical bench, and then used to measure the fluxes. Other parameters are measured in order to estimate the flow according to the following equation:

The equation for *F_c_*is:

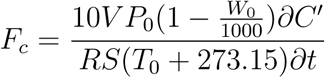

Where: *F_c_*is the soil CO_2_ efflux rate (*µ*mol m*^−^*^2^ s*^−^*^1^), *V* is volume (cm^3^), *P*_0_ is the initial pressure (kPa), *W*_0_ is the initial water vapor mole fraction (mmol mol*^−^*^1^), *S* is soil surface area (cm^2^), *T*_0_ is initial air temperature (°C), and 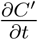 is the initial rate of change in water-corrected CO_2_ mole fraction (*µ*mol mol*^−^*^1^).

The LI8100A analyzer is connected to the SoilFluxPro™ program, which simplifies the management of large datasets and provides post-processing tools to obtain soil gas flux results. This software is specifically developed for and included with LI-COR soil gas flux systems.

The measurements of soil respiration were done randomly on different days at different hours to eliminate bias given by the same weather conditions.

Prior to each measurement, the ground vegetation was cut to avoid a bias due to the respiration of the aerial parts of the plants. Then the chamber, the analyzer control unit, and the temperature and relative humidity (RH) probes were connected to each other. All these components were then linked to a computer running the LI-8100 Automated Soil CO_2_ Flux System software, which was used to acquire measurement parameters. Once the area was ready a PVC collar was placed 3 cm deep into the soil, and a closed transparent chamber was positioned above it to ensure the chamber was airtight. The chamber was then attached to the LICOR Analyzer Control Unit, and the measurement of soil CO_2_ flux started once the chamber reached 50°C. Each measurement lasted 2 minutes, with 30 seconds of pre-purging and 30 seconds of post-purging between each measurement. Three replicates were carried out for each measurement. To ensure uniform microclimatic conditions inside and outside the chamber, the time of the measurement needs to be short. Following the fieldwork, the soil samples were analyzed in the laboratory facilities of the University of Lausanne. Here, analyses took place from September 2023 to December 2023.

### 2.3 Laboratory work

The laboratory methodology follows the protocols of the University of Lausanne. Analysis, including a concise background summary, will be provided here. Please refer to the protocols on the University website for detailed procedures (Soil and Sediment - UNIL). All the samples were initially sieved to 2mm using a square-mesh sieve, and the mass of coarse fragments was reported. Soil is typically sieved under 2mm to ensure a standardized particle size for analysis. Particles smaller than 2mm are classified as fine soil particles, which include sand (0.05 - 2 mm), silt (0.002 - 0.05 mm), and clay (<0.002 mm). These fine particles play a crucial role in soil structure, nutrient retention, and water-holding capacity. By sieving the soil to exclude particles larger than 2mm, which mainly consist of coarse sand, gravel, stones, and organic debris, it is possible to focus on the fine soil fraction that is more involved in the chemical and biological processes (Carter & Gregorich, 2007).

The standard soil analyses were then carried out on sieved soil. For each soil analysis, between 10 and 20% of the initial number of samples were duplicated in order to assess the relative error and reliability of the measurements. An error of up to 10% has been accepted.

**pH** pH determination in a soil suspension provides an estimate of the acidity of a solution in equilibrium with the soil. In this analysis, the acidity is measured in a soil suspension using a combined electrode pH-meter. The ratio between soil:solution influences the results of soil pH measured in water, it is thus important to use a constant ratio and report it alongside the results. The pH of sieved soil was measured in deionized water in accordance with AFNOR standard NF X-31-103, 1998, using a soil/solution ratio of 1:2.5.

### Loss Of Ignition and Soil Organic Matter

To obtain an estimate the soil organic matter (SOM) the method that has been used is the loss of ignition (LOI). Throughout this process, the soil samples are combusted at a temperature high enough to burn off organic material, which then volatilizes as CO_2_ and other compounds, but not so high as to decompose carbonates. The mass loss represents the organic matter content.

### Granulometry

One of the parameters of the description of alpine soil is the characterization of the particle size distribution of mineral particles <2 mm. The laser diffraction particle sizing method is employed to gain insights into granulometry. First, the organic matter was removed by adding H_2_O_2_ 10% and 35% while buffering the pH between 6.5 and 7.5 with 0.1M NaOH. This operation was repeated for several days until the reaction stopped, indicating the complete removal of organic matter. Afterward, the particles were dispersed by adding 1mL of 40g/L Na hexametaphosphate solution, then stirred to keep the elementary particles in suspension for laser analysis. Finally, a Beckmann Coulter granulometer was used to measure particle size distribution between 0.01 *µ*m and 2 mm.

### Organic carbon and nitrogen content (CHN)

The analysis of organic carbon and nitrogen content (CHN) in soil involves several steps to quantify these elements. Before detecting soil organic carbon, acid fumigation is carried out to remove soil carbonate. This process involves exposing the soil sample to acid vapors, which react with and remove the carbonate compounds present. This step is crucial to ensure that the measured carbon content reflects the organic carbon content of the soil without interference from inorganic carbonates. Following acid fumigation, the soil sample is prepared for CHN analysis, often through combustion or other chemical methods. Organic carbon is a key indicator of soil health, it regulates terrestrial ecosystem functioning, provides diverse energy sources for soil microorganisms, governs soil structure, and regulates the availability of organically bound nutrients (Paul & Frey, 2023).

### Available nutrients and total elements

To estimate available nutrients in soil the method used for the extraction is the Mehlich methodology (Mehlich, 1984). Soil samples undergo ICP (Inductively Coupled Plasma) spectroscopy, a technique that uses an inductively coupled plasma to ionize the sample and to identify the target elements (Mylavarapu et al., 2014). The nutrients obtained here are P, K, Ca, Mg, and Mn.

Finally, to observe the distribution of the inorganic forms of various elements, the X-ray fluorescence (XRF) was carried out. This method allows for the detection of major elements in soil, determining the proportion of oxides of the following elements: Si, Al, Fe, K, Mg, Ca, Na, Ti, Mn, and P. In this analysis, fused beads have been prepared using dilithium tetraborate (Li_2_B_4_O_7_) and then burned at 1200°C, to obtain a molten glass pellet. The glass pellet was then subjected to XRF analysis to determine the sample proportions of the various oxides.

Studying the concentration of available elements in soil within the context of glacier retreat is of foremost importance as their availability in soil influences the colonization and establishment of vegetation in newly deglaciated areas.

### 2.4 Data processing and statistical analysis

Statistical analyses were performed on these using RStudio software (version 4.3.1) to answer the research questions and test the hypotheses of this work. First, using Excel all data were organized and grouped depending on the data concerned: vegetation, soil properties, and soil respiration. In RStudio the initial step involved ensuring the distribution of data frequencies through histograms and descriptive statistics. These histograms were used to visualize the data distribution for each dependent variable, whereas residuals and fitted values were used to determine the normality of the data. In cases where the data did not conform to a normal distribution, the respective variables were transformed using suitable methods, such as logarithmic transformations to achieve a more normal distribution.

In this study, we employed linear models (i.e., lm) to assess the relationship between our dependent variables and the non-dependent variable. The dependent variables, also known as response variables, considered in this analysis included the soil parameters (e.g., soil organic matter, carbon and nitrogen content, pH, soil respiration), and vegetation data (e.g., richness, diversity) (see Figure 9). The non-dependent variables, also known as predictors or independent variables, consisted of categorical (Stage) and numerical (Years since deglaciation) variables, used to explain changes in our response variables. Using both predictors enabled the creation of similar but complementary models. Running statistical analyses with the categorical predictor "Stage" helps determine whether the stages are distinct from each other, while using the linear predictor "Years" allows for the observation of the distribution trend of the data along the ice-free areas. Using both predictors allows the analysis to be strengthened. The results obtained are qualitatively similar, hence, we will only consider the results obtained with one predictor, Years.

**Figure 9:**
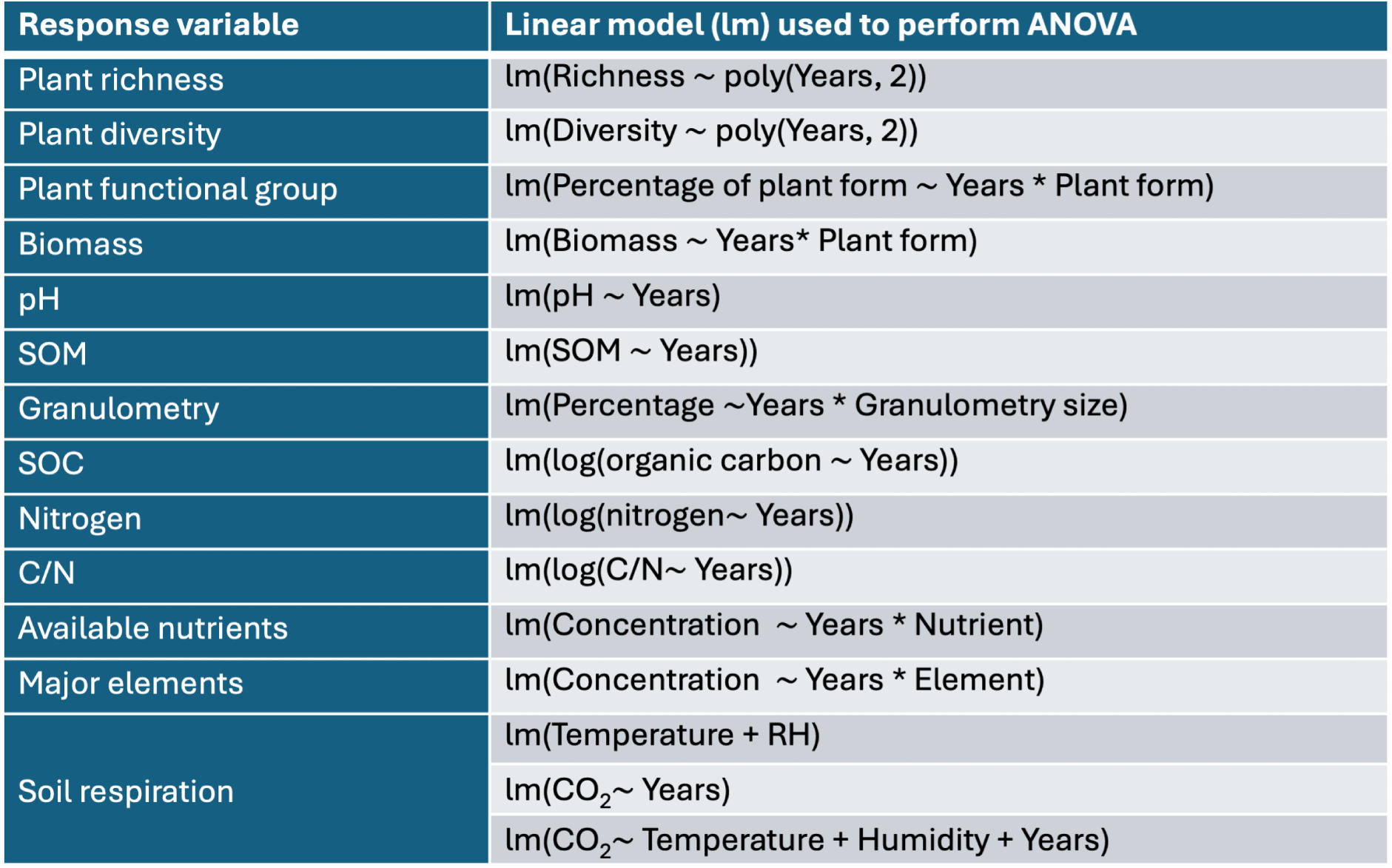
The analysis of variance (ANOVA) was performed according to the following model, for each response variable we wanted to investigate

For every model analyzed we performed a hierarchical ANOVA (Analysis of Variance). Hierarchical ANOVA is a statistical technique used to analyze the variance in a dependent variable by grouping the independent variables into different levels. We evaluated three separate models for each parameter under investigation and chose the most suitable one based on the F-value and p-value derived from the ANOVA. The three models are as follows:

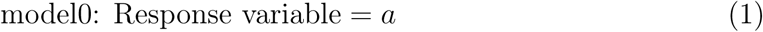

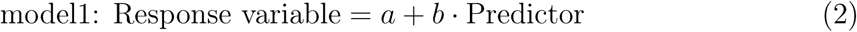

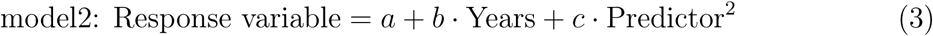

Model0 (1) represents a simple linear regression model, with the response variable explained by a constant, model1 (2) represents a simple linear regression model, with the response variable explained by a predictor, finally model2 (3) represents a second-degree polynomial equation, with the response variable explained by a predictor with a polynomial transformation.

Eventually, the models that best fit the data are shown in the following figure 9:

The criteria used to assess the model’s suitability and determine the optimal fit included the F-value and the associated p-value. The F-value is used to test whether the variation explained by the model is statistically different from the unexplained variation. The p-value typically below a predetermined significance level such as 0.05, suggests that the observed data are unlikely to have occurred under the assumption of no effect or no difference. Thus, by examining these statistical metrics, it is possible to evaluate the data distribution suitability and to choose the best-fitting model. Additionally, using the Akaike Information Criterion (AIC) it was possible to evaluate how well each model was fitting the data. Preference was given to the model with the lowest AIC, indicating a better balance between data fit and model complexity. After selecting the best-fitting model, a Type-II ANOVA was conducted on the linear regression model to assess if the predictor statistically explained the variation in the response variable.

Moreover, the Tukey’s Honestly Significant Difference (HSD) test was employed as a critical tool for analyzing the effects of glacier retreat on our dependent variable. The Tukey’s test allows for comprehensive pairwise comparisons within the different "Years" or "Stage". By identifying statistically significant differences in the response variable over the years since deglaciation, it’s possible to gain a deeper understanding of how vegetation, soil, and soil respiration respond to varying degrees of glacier retreat. Ultimately, the data were graphically represented through boxplots and/or scatter plots, providing a visual representation of the findings.

Principal Component Analysis (PCA) was conducted to explore the multivariate relationships among the variables in our dataset. A PCA is a statistical technique used to reduce the dimensionality of data while maintaining as much variance as possible. The results of PCA were visualized using the fviz_pca_var function from the factoextra package in R. Additionally, to examine the interactions among soil respiration, soil characteristics, vegetation, and glacier retreat, a Structural Equation Analysis (SEM) was conducted using the lavaan package in R. SEM was performed on the small dataset size, thus represents a simplified path analysis for a limited number of variables.

Variables selected for this model are years since deglaciation, plant diversity, soil organic carbon, SOM, biomass, and soil respiration (Soil CO2 flux), and are represented by the model (28) as the following equations:

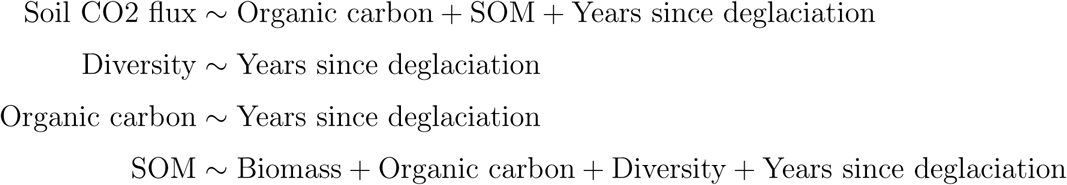

## 3. Results

### 3.1 Vegetation

#### Plant richness and plant diversity

Glacier retreat influences plant richness (F-value = 34.76, p < 0.05). As shown in Figure 10, plant richness (i.e., the number of species) increases along the chronosequence of glacier retreat, reaching a maximum at stage 2, the intermediate stage, with a median number of plant species at 38. The lowest richness value is observed in stage 0, representing the most recently ice-free area, with a median richness of 11 species. Following the peak, species richness decreases in the later stages, reaching the second-to-last lowest values in stage 4, with a median of 12 species. Richness increased by 30% from stage 0 to stage 2. Significant statistical differences (p <0.05) in species richness among the different stages were found among stages 0 and 2, 0 and 3, 2 and 4, and 3 and 4.

**Figure 10:**
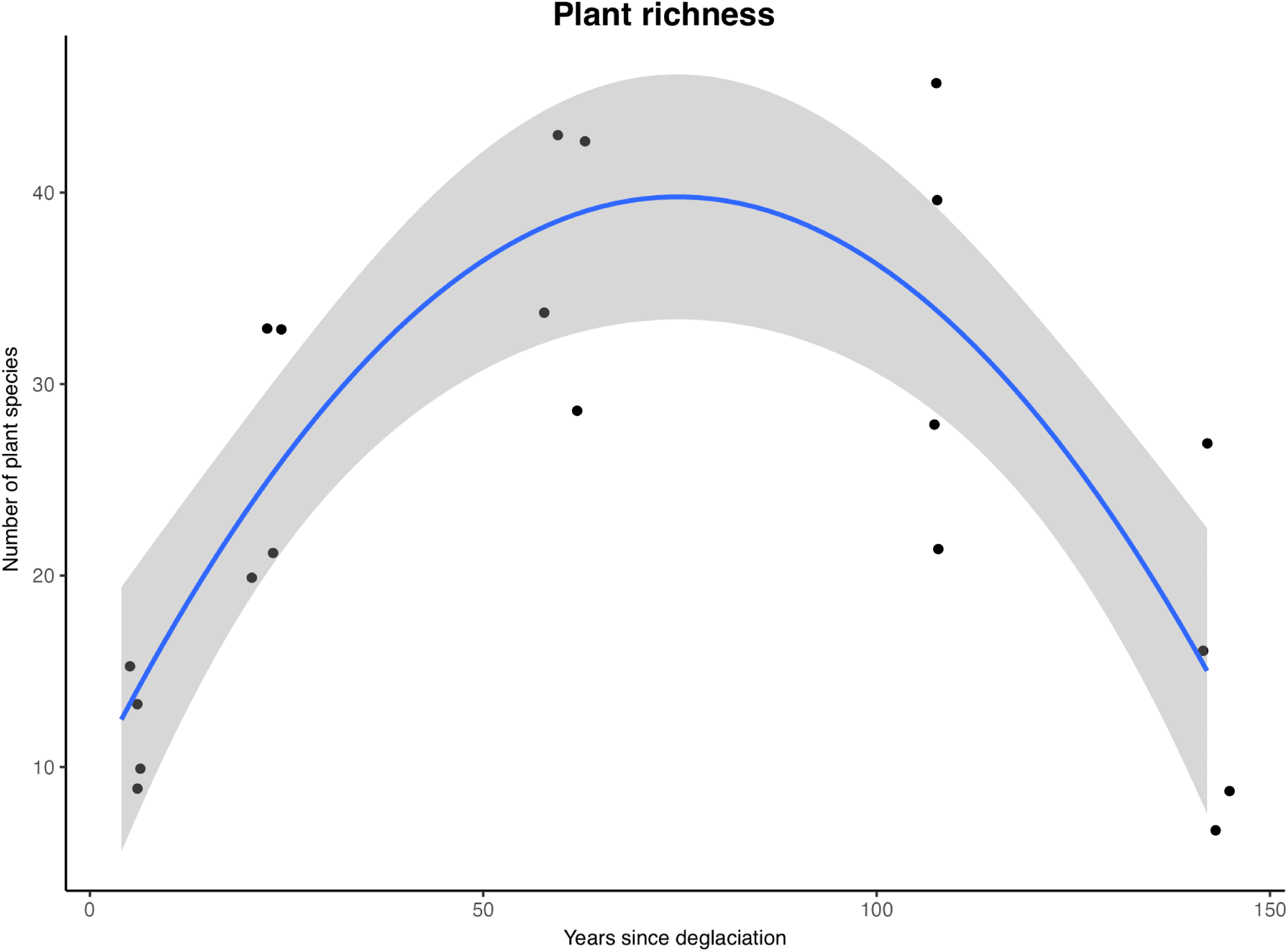
Boxplot of plant richness according to glacier retreat. Richness corresponds to the sum of species present in each plot and glacier retreat is represented by years since deglaciation and stages.

The same trend of plant richness is observed in plant diversity (i.e., Shannon Index), which is also influenced by glacier retreat (F-value = 31.26, p <0.05). Plant diversity increases along the primary succession (Figure 11) reaching a peak at the intermediate stage and then decreasing. Diversity at stages 0 and 1 is respectively 2.3 and 2.4. It reaches the maximum value at stage 2 with a median of 3, then declines to 2.9 in stage 3. Finally, it falls to the lowest value within stage 4, recorded at 1.7. Statistically significant differences (p <0.05) in plant diversity are found between stages 0 and 2, 1 and 2, 2 and 4, and eventually 3 and 4. All the other stages were not statistically different from each other.

**Figure 11:**
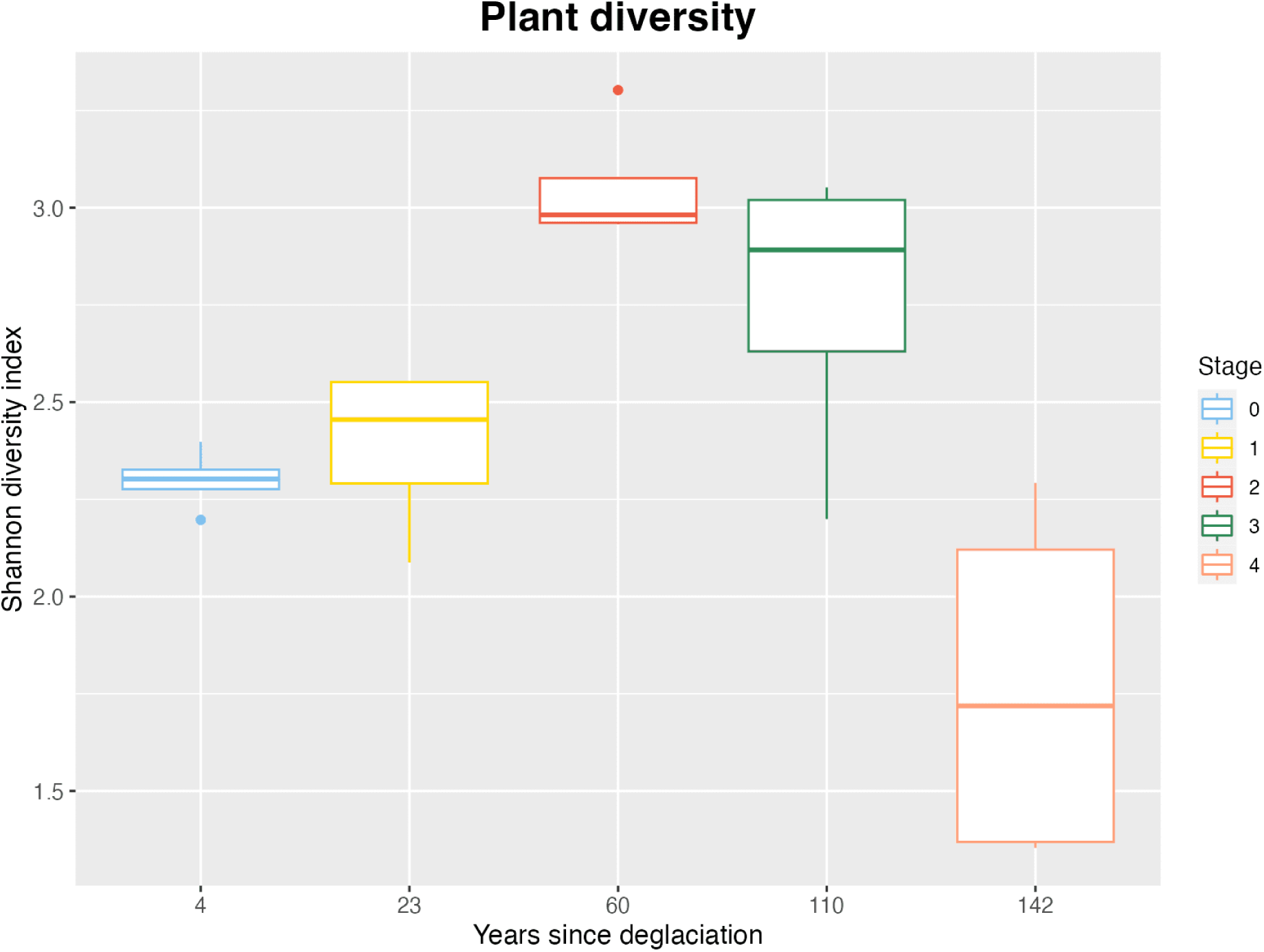
Boxplot of plant diversity according to glacier retreat. Diversity is calculated using the Shannon Index for each plot and glacier retreat is represented by years since deglaciation and stages.

### Plant functional groups

Glacier retreat influences the proportion of different plant groups occurring along the chronosequence (Figure 12) (F-value = 64.53, p <0.05). Statistics confirm that glacier retreat influences proportion of forbs (t-value = 15.22, p-value <0.05), graminoids (t-value = 2.82, p < 0.05), shrubs (t-value = 5.92, p < 0.05), and trees (t-value = 4.35, p < 0.05).

**Figure 12:**
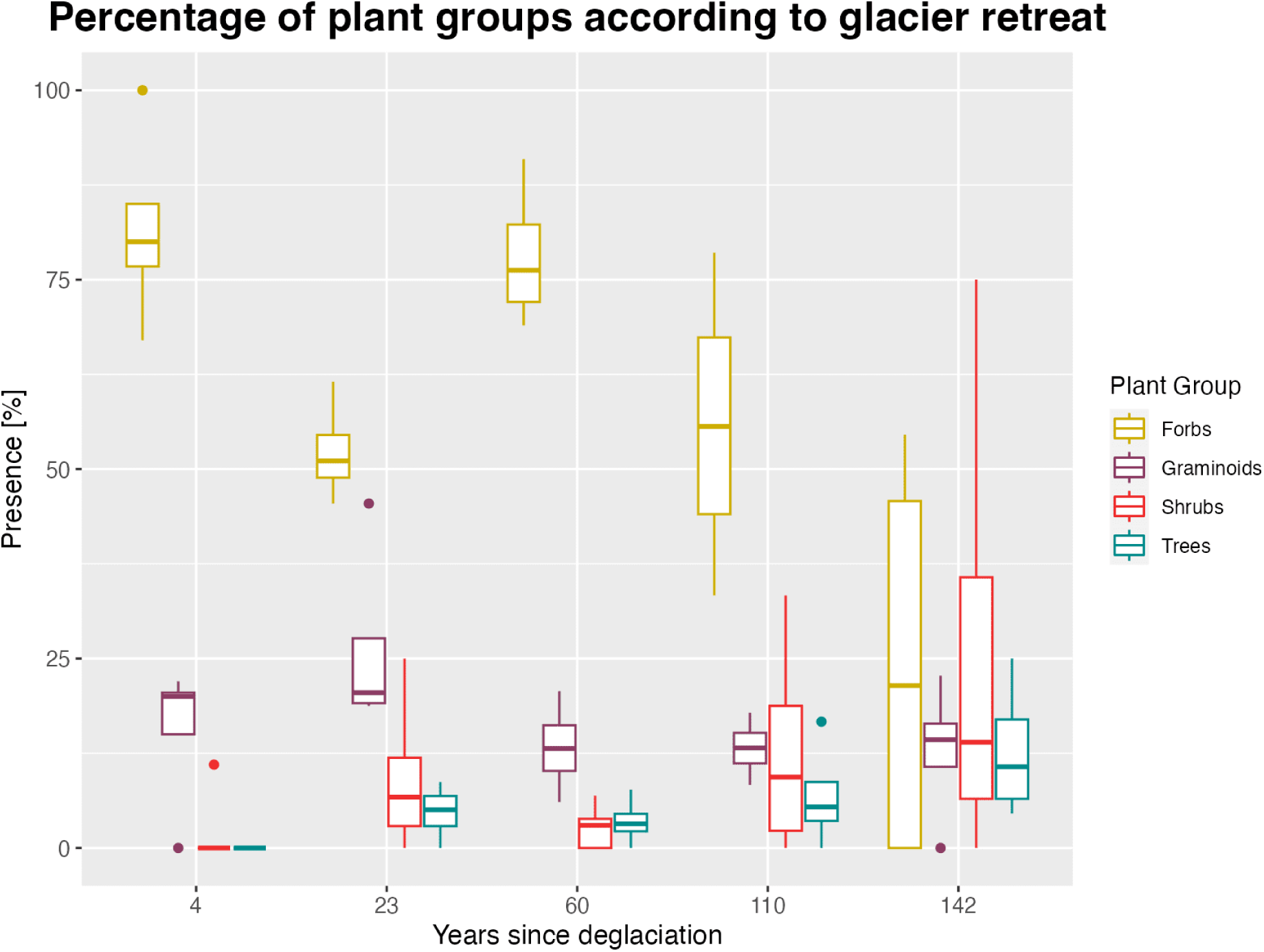
Representation of plant functional groups along the gradient of glacier retreat. Each data point represents the occurrence of a specific plant group within a plot. Glacier retreat is represented by years since deglaciation corresponding to 4, 23, 60, 110 and 142 years associated with stages 0, 1, 2, 3 and 4 respectively.

Forbs and graminoids have the highest presence in the early and intermediate stages, until stage 3, where their presence falls to the detriment of shrubs and trees. The median value for the proportions of forbs is 80% in stage 0, 51% in stage 1, and then reaches the maximum of 76% in stage 2, while for graminoids the presence is around 20% for both stage 0 and 1 and then decreases to 13% in stage 2. Stage 3 is also dominated by forbs (median value of 55%), before decreasing in stage 4 (median value of 21%). From stage 3, trees and shrubs become the most present plant form. Regarding shrubs and trees, an early-stage willow (*Salix sp.*) has been observed already at stage 0, where trees are still absent. Trees and shrubs have a low presence in stage 1 (16% and 18% respectively), but they become progressively more abundant as the primary succession advances.

### Biomass

Figure 13 shows the trend of biomass (i.e., carbon stock in aboveground vegetation) along the primary succession. The change in biomass following glacier retreat is statistically significant (F-value = 19.85, p <0.05). Only stage 4 is statistically significantly different from the other stages (p <0.05). At the pioneer stage, 4 years since deglaciation, there’s no carbon accumulated in aboveground vegetation. As terrain age increases, the carbon stock in aboveground biomass accumulates mostly in trees, whereas shrubs and graminoids only account for a small part of it. Estimated total above-ground biomass (AGBest) at the pioneer stage is 0, then it increases to 300 kg already in stage 1, because of the shrubs and woody pioneer plant species and reaches 14 346 kg in the late successional stage.

**Figure 13:**
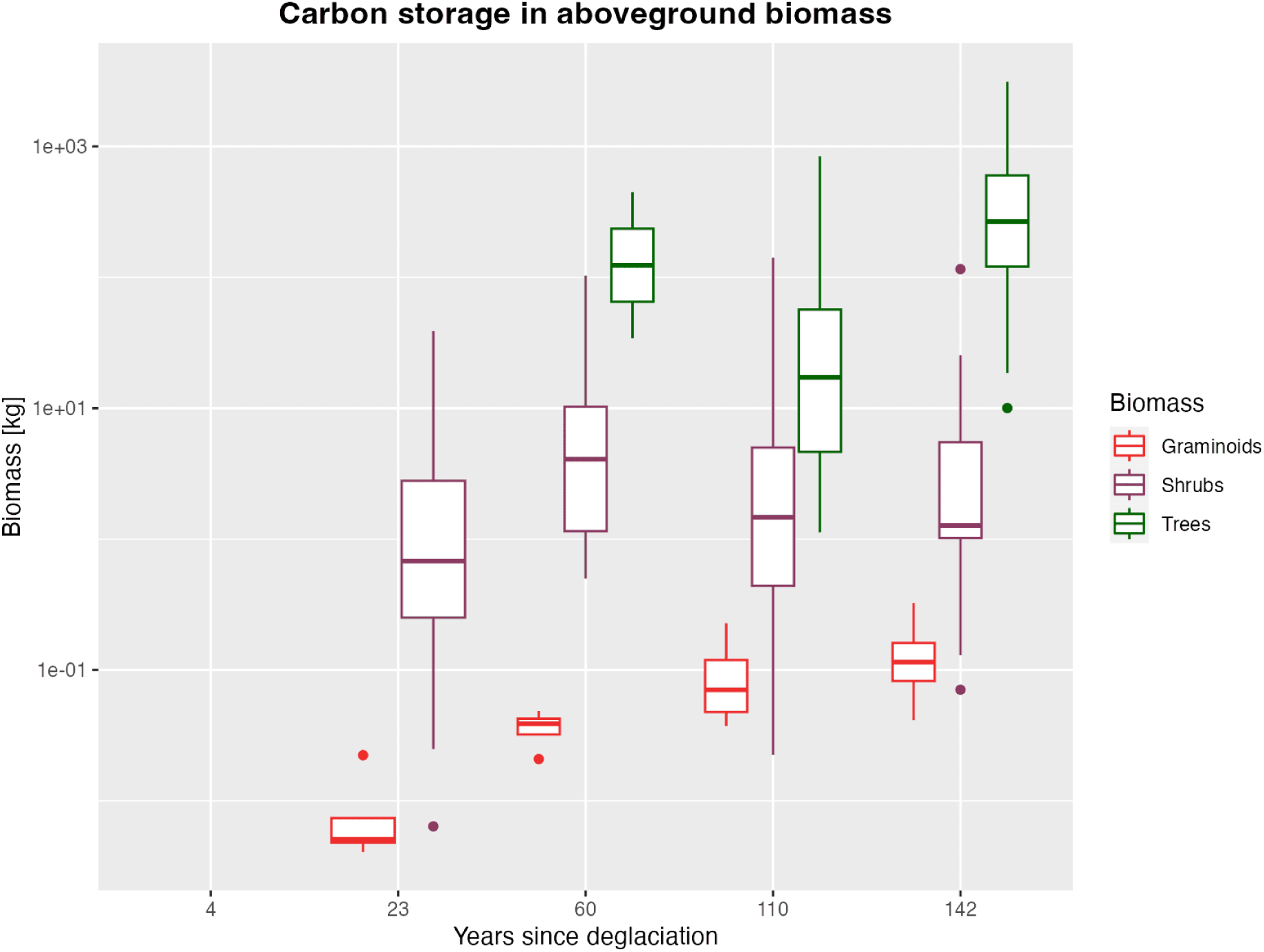
Estimate of carbon storage in aboveground biomass. Biomass is represented in logarithmic scale. Glacier retreat is represented by years since deglaciation corresponding to 4, 23, 60, 110 and 142 years associated with stages 0, 1, 2, 3 and 4 respectively.

### 3.2 Soil development

#### pH

Figure 14 illustrates a decrease in soil pH following glacier retreat (F-value = 231.65, p<0.05). The pH values are highest at stage 0, with a median value of 6.9 indicating neutral conditions. pH decreases in the following stages to 6.6, 5.9, 5.26, and reaches its lowest value of 4.3 at stage 4, reflecting increasingly acidic conditions.

**Figure 14:**
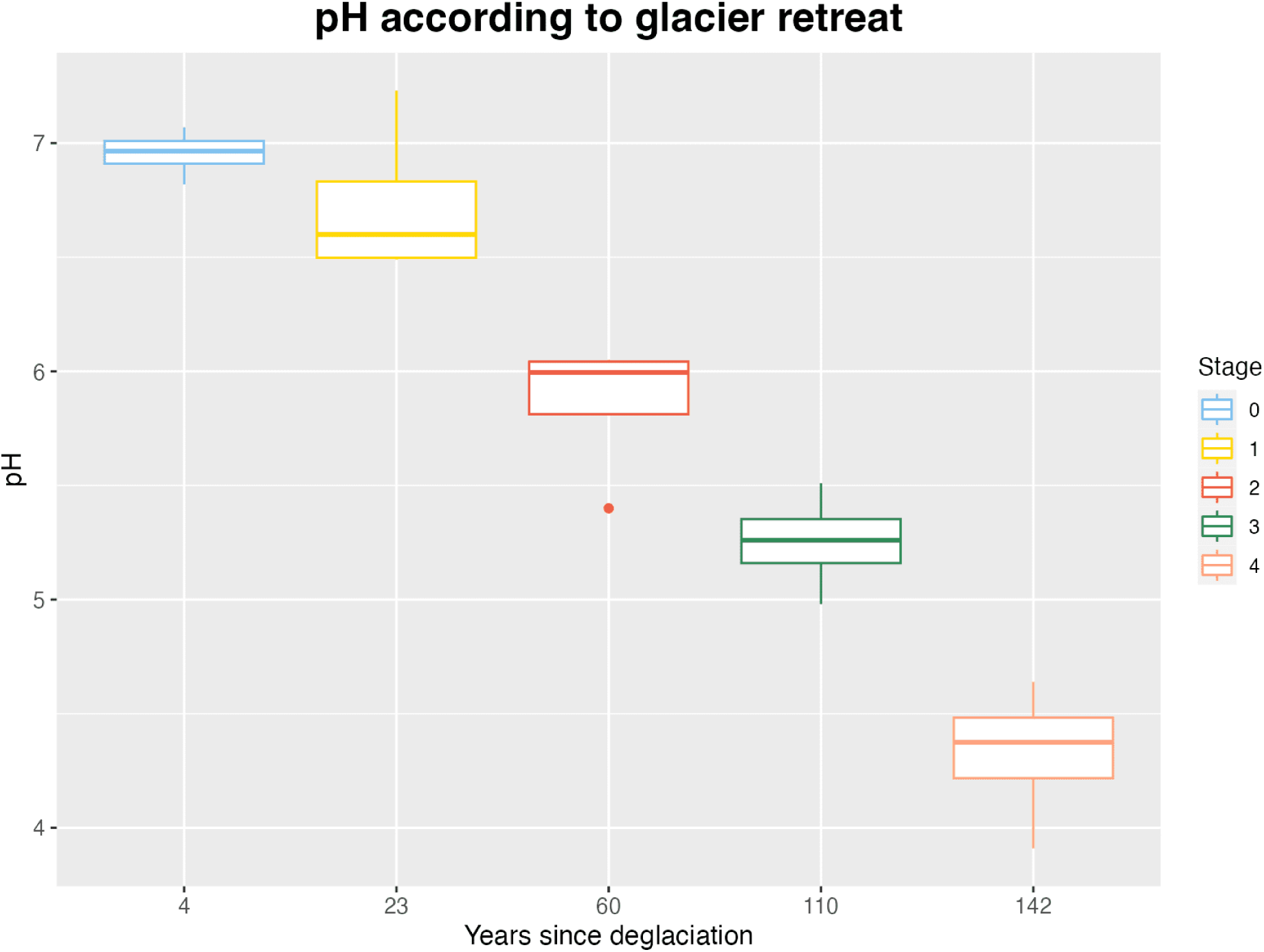
pH according to glacier retreat. Glacier retreat is represented by years since deglaciation and colors correspond to the respective stages.

According to the Tukey test, all stages are significantly different from each other (p<0.05).

### Soil Organic Matter

Figure 15 illustrates an increase in soil organic matter (SOM) following glacier retreat. Glacier retreat has a significant impact on SOM (F-value = 21.13, p <0.05). The SOM values are very similar in the early stage (median values of 0.11% and 0.65% for stages 0 and 1 respectively) and in the intermediate stage (4.76% and 3.88% for stages 2 and 3 respectively). The SOM content increases steeply in the late successional stage, reaching a value of 14.1%. According to the Tukey test, only stage 4 is significantly different from the other stages (p<0.05), whereas the other stages are not significantly different from each other (p >0.05).

**Figure 15:**
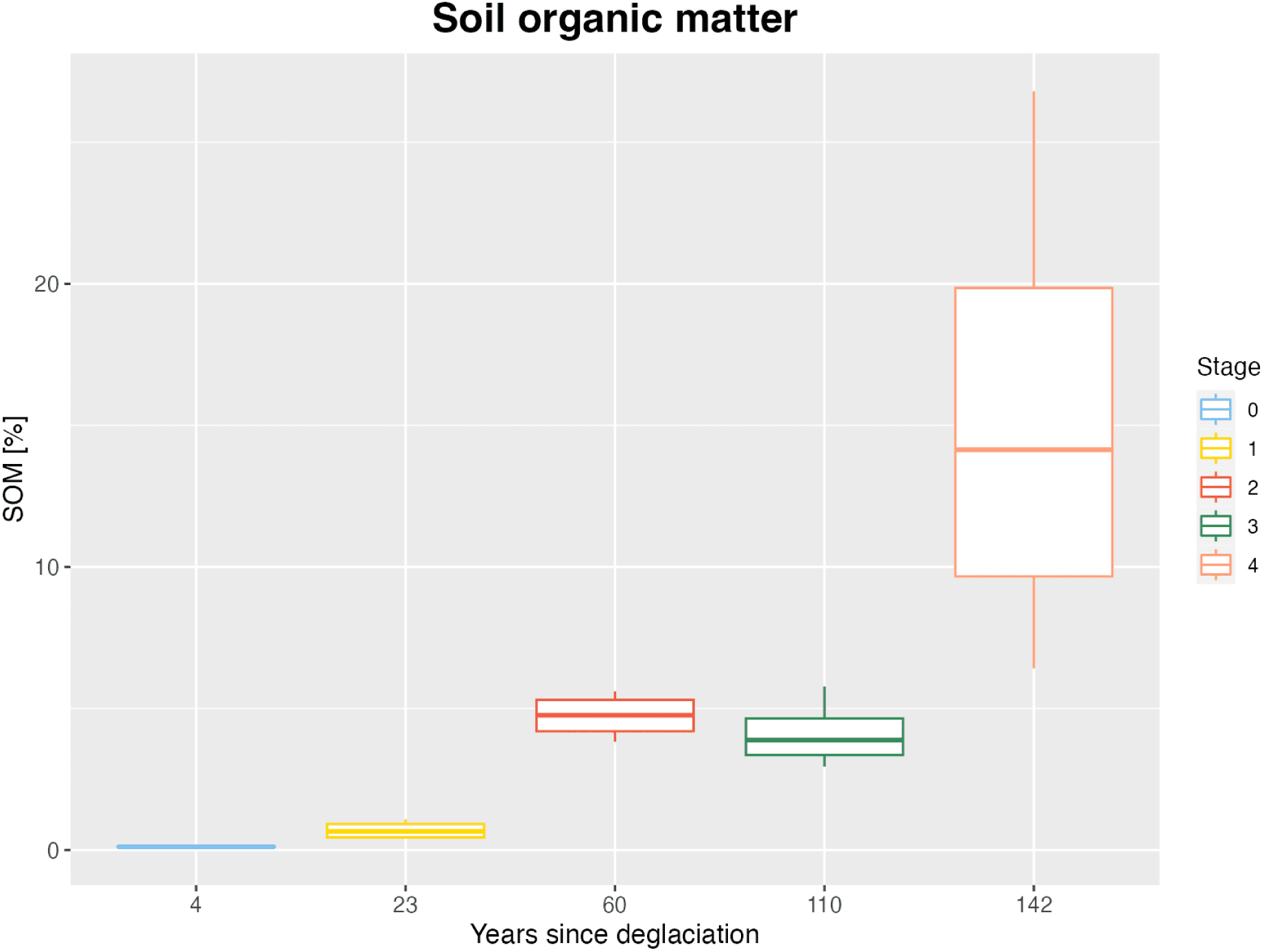
Boxplot of soil organic matter (SOM) according to glacier retreat. Glacier retreat is represented by years since deglaciation and colors correspond to the respective stages.

### Granulometry

The texture is affected by glacier retreat (F-value = 6.74, p <0.05). As Figure 16 shows, the percentage of sand remains the most abundant all over the stage (with an overall average of 75%), whereas the clay represents the least abundant (overall of 2%). The percentage of silt increases following glacier retreat, from an average of 2% in stage 0 to an average of 36% in the late successional stage.

**Figure 16:**
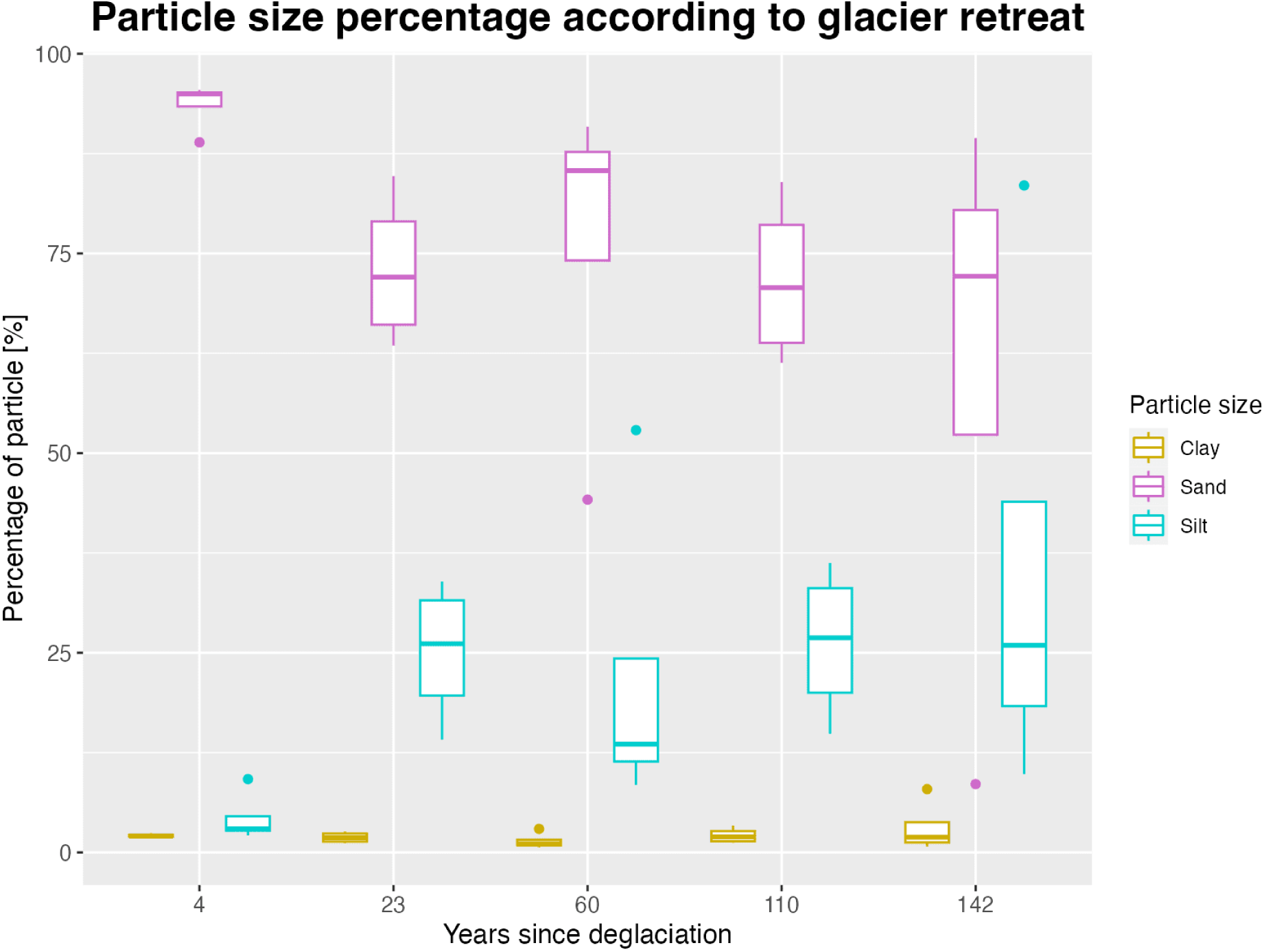
Texture

### Soil organic carbon and total nitrogen

The content of soil organic carbon (SOC), total nitrogen (Ntot), and their ratio (C/N ratio) is impacted by glacier retreat, as indicated by the statistical analysis (respectively, F-value = 74.41, p <0.05; F-value = 24.24, p <0.05; F-value = 72.87, p <0.05).

Regarding organic carbon, as Figure 17 shows, from almost a null amount in stage 0 (median value of 0.0344 %), organic carbon content increases rapidly, reaching the maximum value of 9.12% at stage 4. The same increasing trend is evident for nitrogen (Figure 18), which from a content of 0.014%, rapidly increases reaching the maximum value of 0.51% in stage 4. The ratio between C and N (C/N) (Figure 19) reflects the carbon and nitrogen trend. The ratio is low at the early stages, accounting for a median value of 2.12, 2.16, and 6.59 respectively for stages 0, 1, and 2. Then the ratio rises, reaching a median value of 8.69 in stage 3, and then the highest in stage 4, with a median value of 17.44. Moreover, the early stages show more homogeneous values and are more aggregated than the later stages, 3 and 4, with more scattered values.

**Figure 17:**
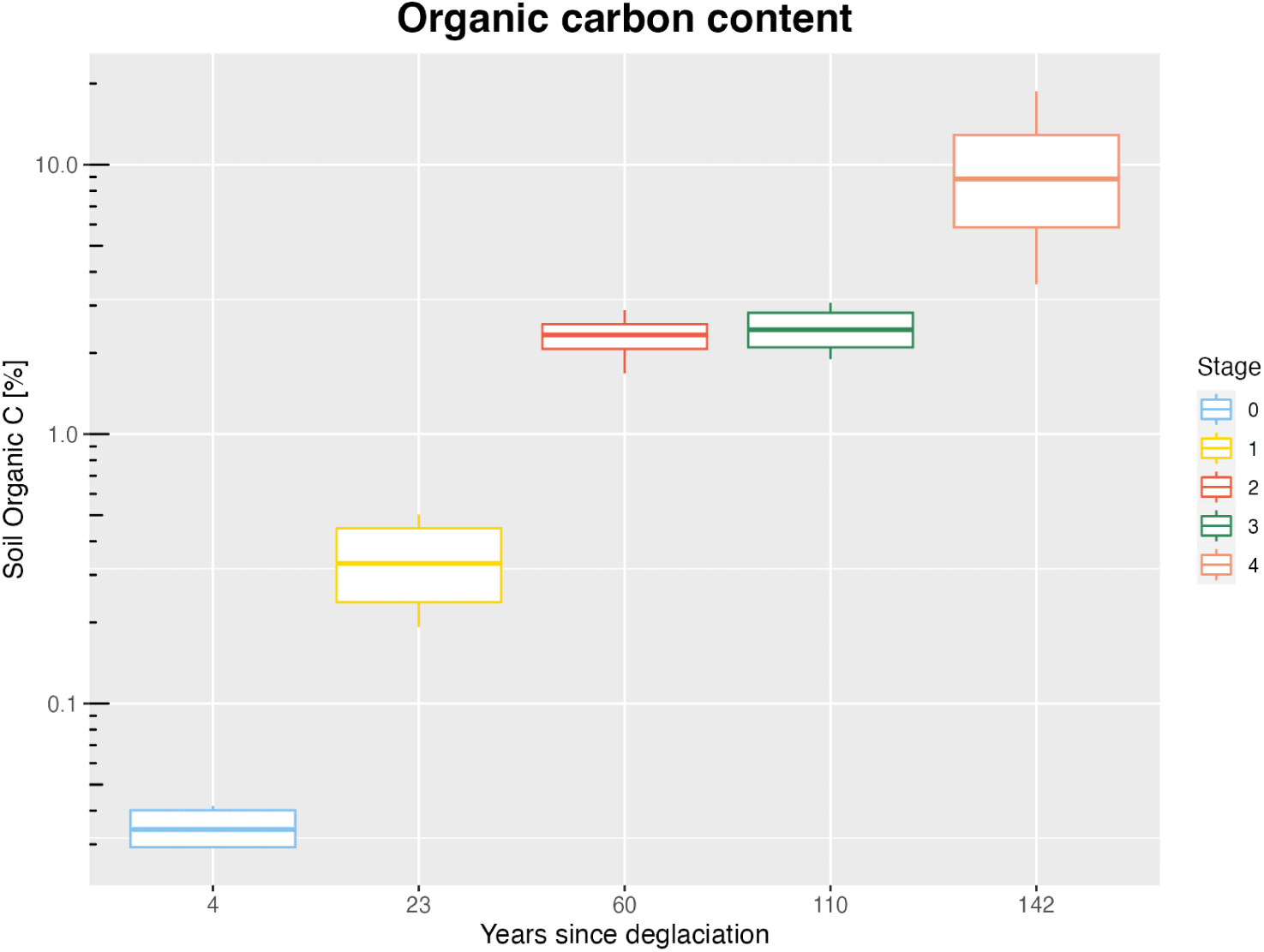
Boxplot of soil organic carbon (SOC) according to glacier retreat. Y scale is log distributed with untransformed percentage of SOC. Glacier retreat is represented by years since deglaciation and stages.

**Figure 18:**
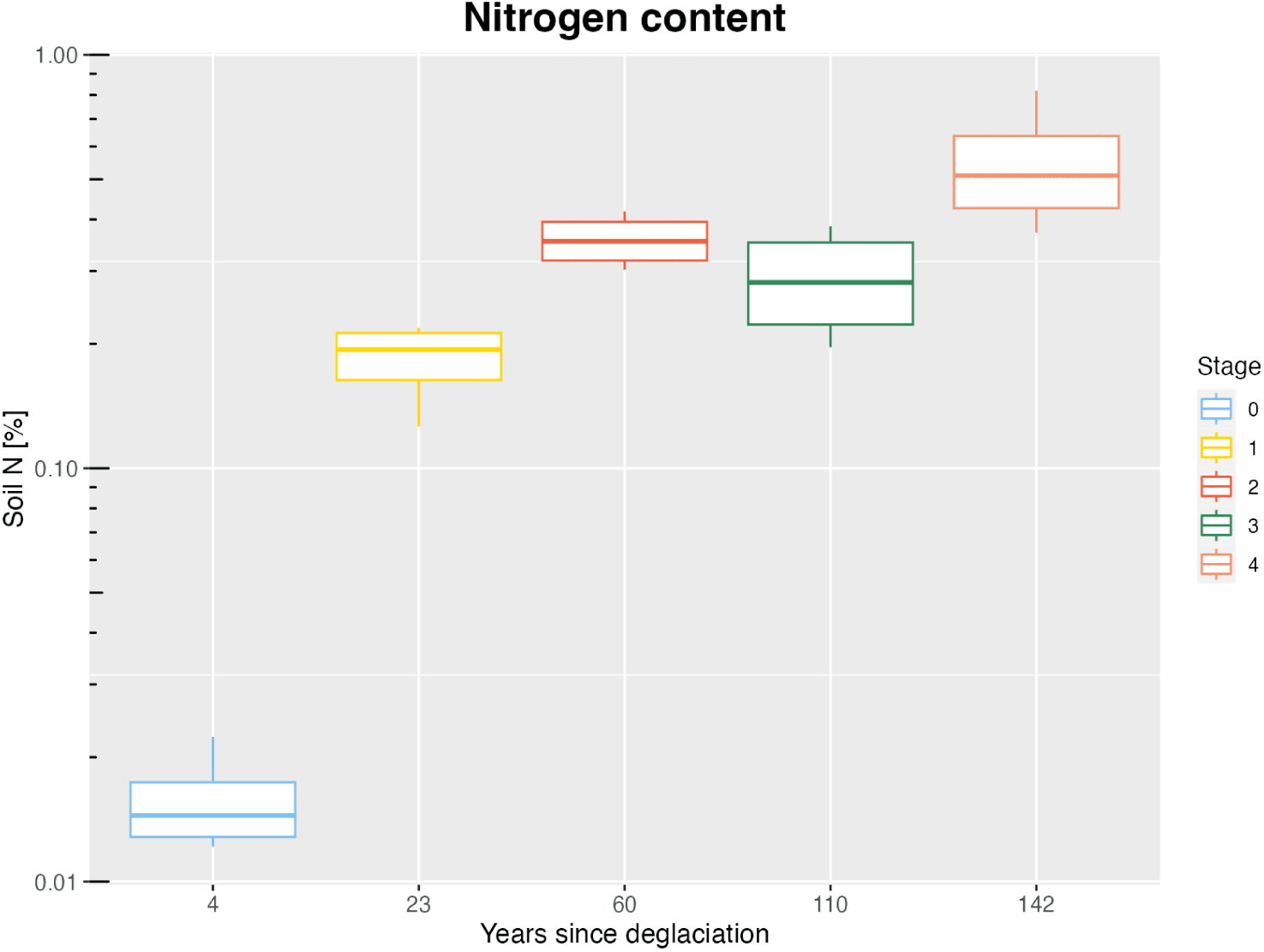
Boxplot of nitrogen content in soil according to glacier retreat. Y scale is log distributed with untransformed percentage of SOC. Glacier retreat is represented by years since deglaciation and stages.

**Figure 19:**
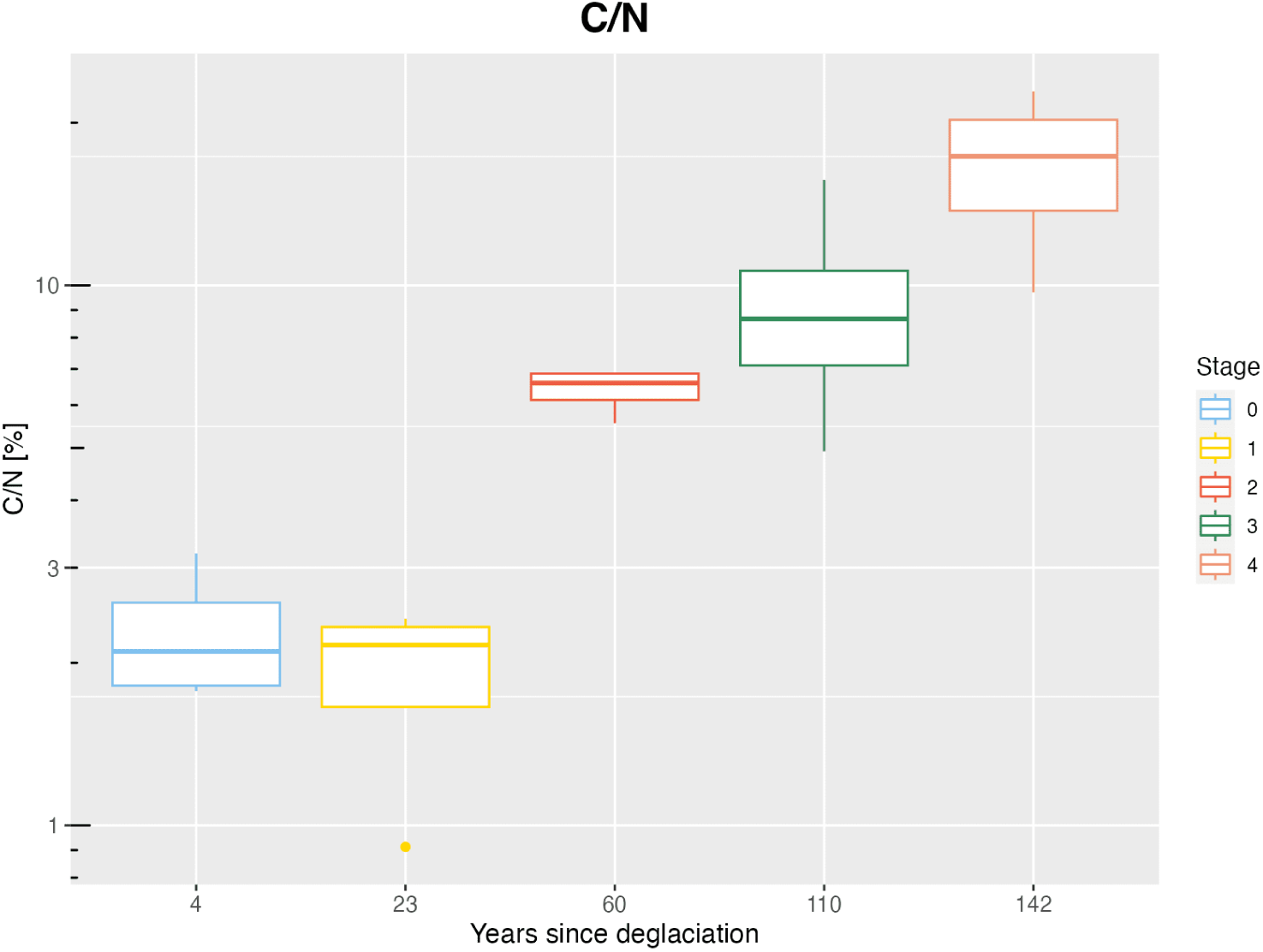
Boxplot of the ratio of C and N (C/N ratio) in soil according to glacier retreat. Y scale is log distributed with untransformed percentage of C/N. Glacier retreat is represented by years since deglaciation and stages.

According to the Tukey test, all stages have statistically significant differences (<0.05) in SOC content, except for stages 2 and 3 (p >0.05). Concerning the nitrogen content only stage 0 and stage 1 are significantly different from the other stages (p <0.05), and all other stages don’t show any statistical difference (p >0.05). Eventually, in the C/N ratio, statistically significant differences are observed among all stages except for stages 0 and 1, 0 and 3, and finally 1 and 3.

### Available nutrients and total elements

The concentration of available elements changes with glacier retreat (F-value = 29.78, p <0.05) (Figure 20). All the measured nutrients have a concentration that increases with glacier retreat. Very low available nutrients characterize stage 0, then the values gradually increase over time following deglaciation. At stage 0 there is no potassium (K) (0 mg/kg of soil), whereas calcium (Ca), phosphorous (P), magnesium (Mg), and manganese (Mn) have very little concentration. With increasing years since deglaciation the concentration of available nutrients in soil increases. Over 140 years calcium increases by a factor of 200 and potassium by a factor of 300. Manganese and phosphorous concentration increase less, those nutrients increase by a factor of 10 and 90 respectively over the succession. Despite the change in concentration among the stages, only stage 4 is statistically significant.

**Figure 20:**
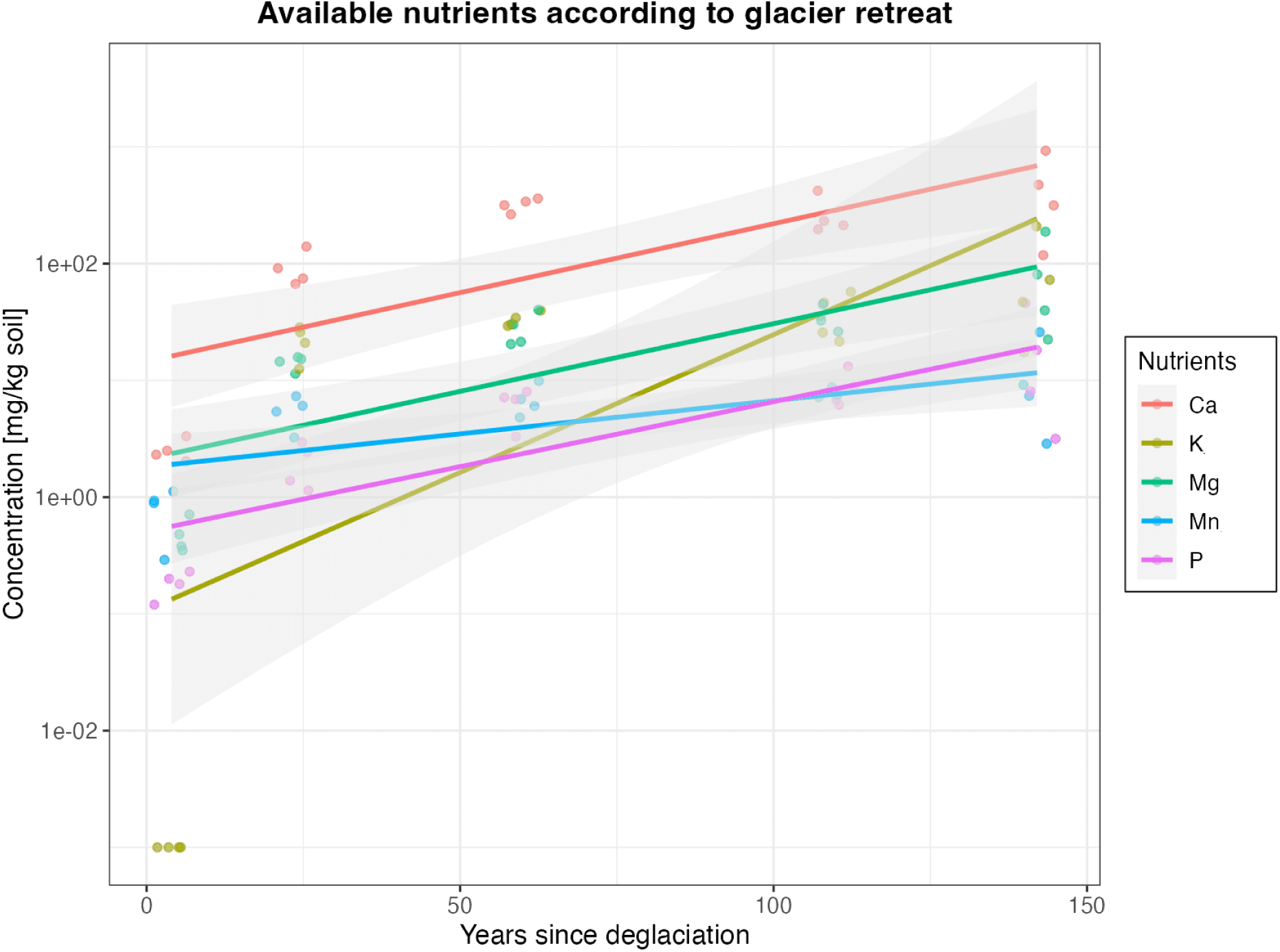
Concentration of available elements according to glacier retreat. Y-scale is a logarithmic scale. Elements are represented as mg of elements per kg of soil and correspond to calcium (Ca), potassium (K), magnesium (Mg), and phosphorus (P). Glacier retreat is represented by years and corresponds to stage 0 (4 years) to stage 4 (142 years).

Comparing the available nutrients with the total proportion of the same elements represented by the percentage of oxides (Figure 21 and 22), the trend is different. Glacier retreat impacts the concentration of total elements in soil (F-value = 2.09, p <0.05). Interestingly, total phosphorus (P_2_O_5_), increases doubling between stages 0 and 4. Total calcium (CaO) and magnesium (MgO) decrease by half between these same stages. Potassium (K_2_O) has also the highest concentration at the pioneer stages, then decreases slightly. Aluminum (Al_2_O_3_), silica (SiO_2_), and sodium (Na_2_O) percentages remain stable over the stages. Manganese (MnO) slightly decreases from stage 0 to stage 1, then increases again.

**Figure 21:**
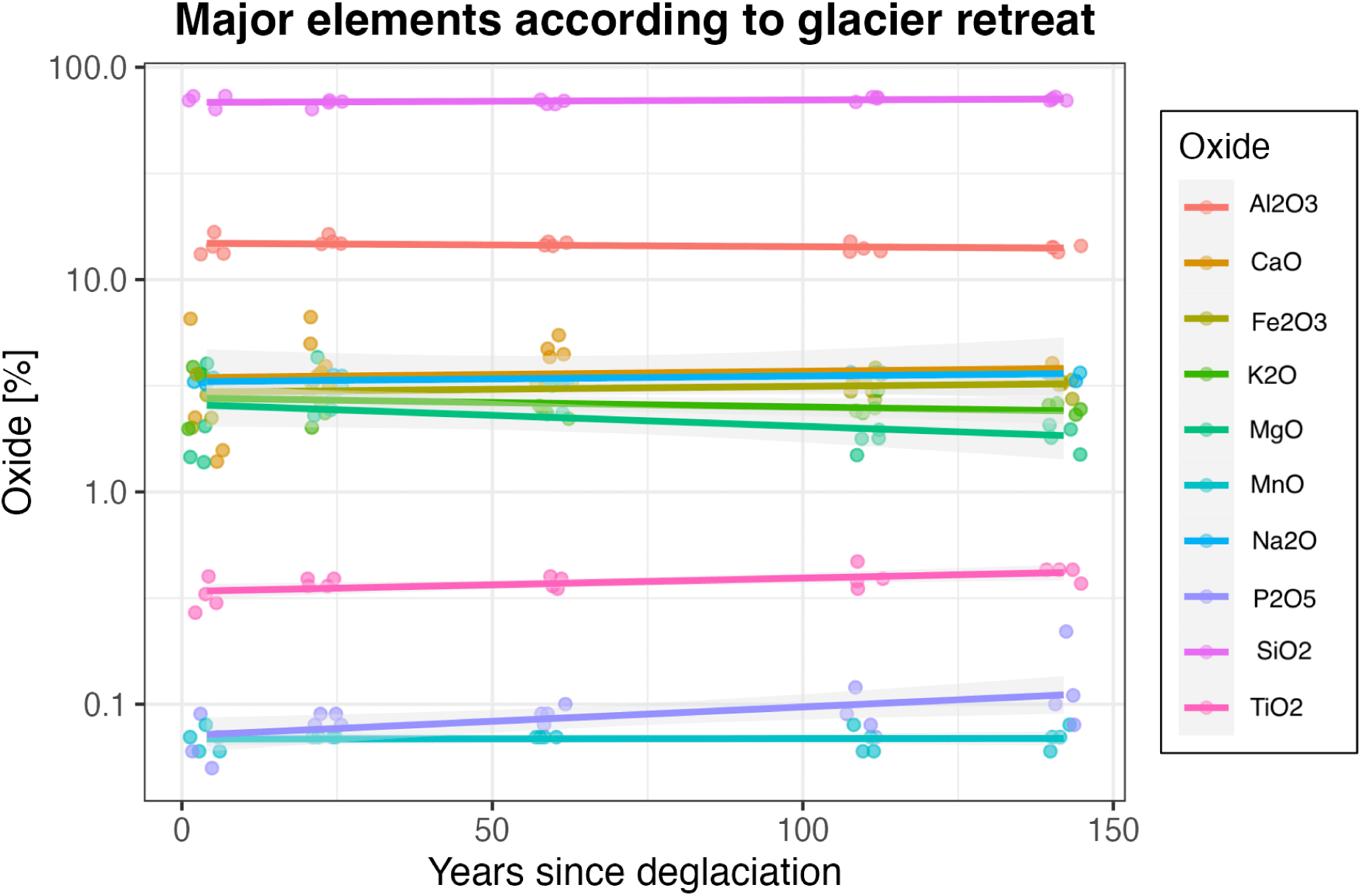
Percentage of major elements representing total elements according to glacier retreat. Y-scale is a logarithmic scale. Elements are represented in the percentage of oxides and correspond to the proportions of each element per stage. This means that in each stage, the sum of all the elements measured here reaches 100%. Elements correspond to Al2O3, CaO, Fe2O3, K2O, MgO, MnO, Na2O, SiO2, P2O5, and TiO2. Glacier retreat is represented by years and corresponds to stage 0 (4 years) to stage 4 (142 years).

**Figure 22:**
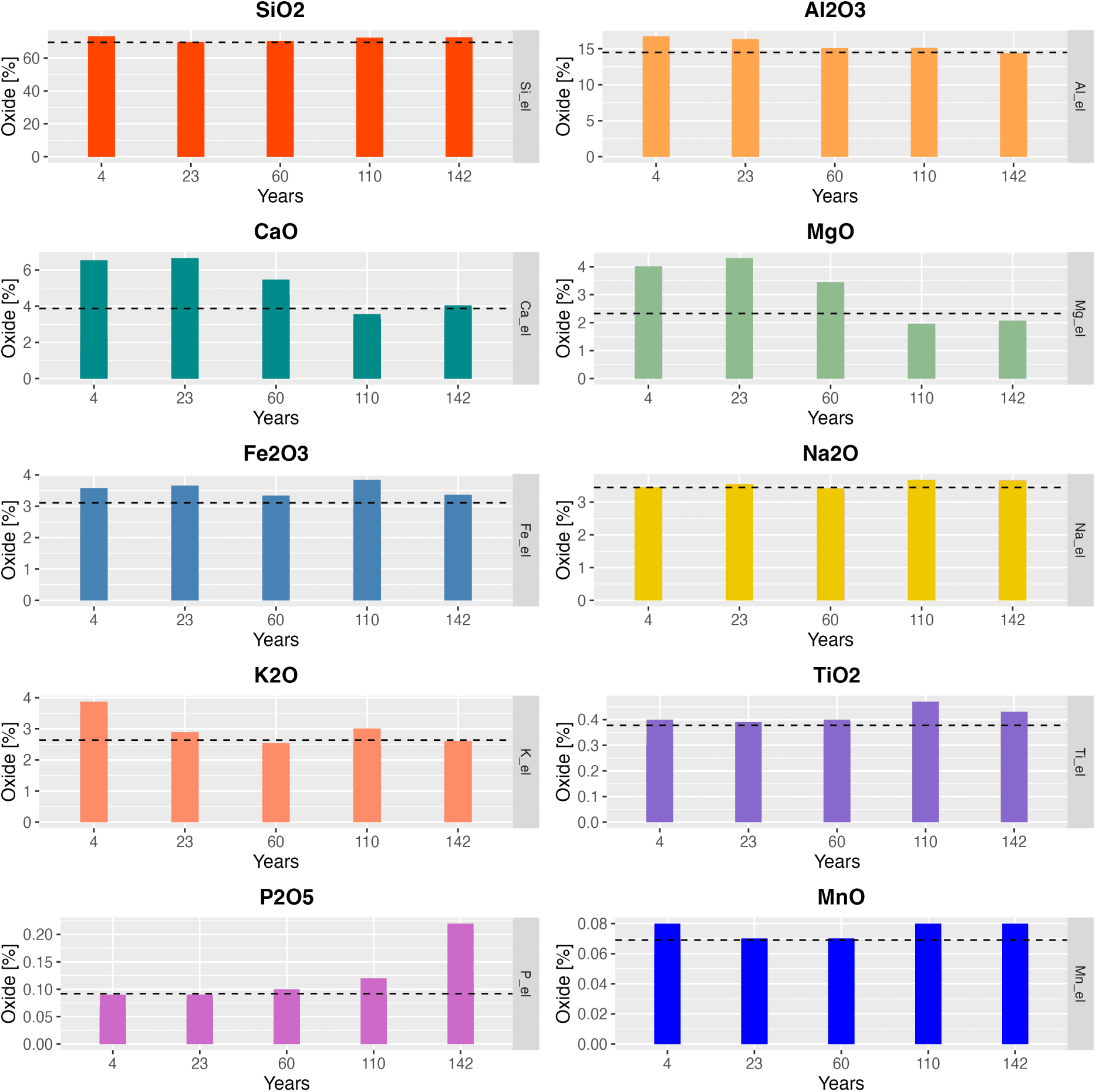
Proportion of major elements representing total elements according to glacier retreat. Elements are represented in percentage of oxides and correspond to the proportions of each element per stage. This means that in each stage, the sum of all the elements measured here reaches 100%. Glacier retreat is represented by years since deglaciation

### 3.3 Soil respiration

Measurements of soil CO_2_ respiration as flux show that the dynamics of respiration vary according to glacier retreat (F value = 73.99, p < 0.05). Due of the challenging placement of the PVC collar, some CO_2_ data, especially in stage 0 where carbon flux is already close to 0, resulted in negative values. Therefore not obtaining bias, data < 0 have been excluded. When testing for statistically significant differences in CO_2_ flux between stages, it was found that all stages are significantly different from each other (p<0.05) except for stages 1 and 4 (p>0.05). The median values of carbon flux start from nearly 0 in stage 0, with a respiration rate of 0.050 *µ* mol m*^−^*^2^ s*^−^*^1^, then slightly increase in stage 1 to 0.752 *µ* mol m*^−^*^2^ s*^−^*^1^. In stage 2, soil respiration rises and reaches the peak with a flux of 1.504 *µ* mol m*^−^*^2^ s*^−^*^1^, followed by a decline to to 1.216 *µ* mol m*^−^*^2^ s*^−^*^1^ in stage 3 and further down to 1.056 *µ,* mol m*^−^*^2^, s*^−^*^1^ in stage 4.

### Temperature and Relative Humidity

Additionally, the effect of temperature and humidity on soil respiration was tested. We first measured how temperature (T) and relative humidity (RH) correlate. Temperature and RH refer to the values obtained by inserting their relative probes into the ground during the fieldwork. Using the Pearson correlation coefficient (r) we found that temperature and RH have a strong negative correlation (r = -0.811). The linear regression model indicates a significant relationship between temperature and RH (F-value = 923.41, p < 0.05), suggesting that changes in temperature are associated with corresponding alterations in relative humidity.

Temperature and relative humidity do not have a significant effect (p >0.05) on soil CO_2_ flux according to the linear model we performed. Moreover, the Pearson correlation coefficient r = 0.034 indicates a very weak and negative correlation between CO_2_ and temperature (Figure 25).

**Figure 23:**
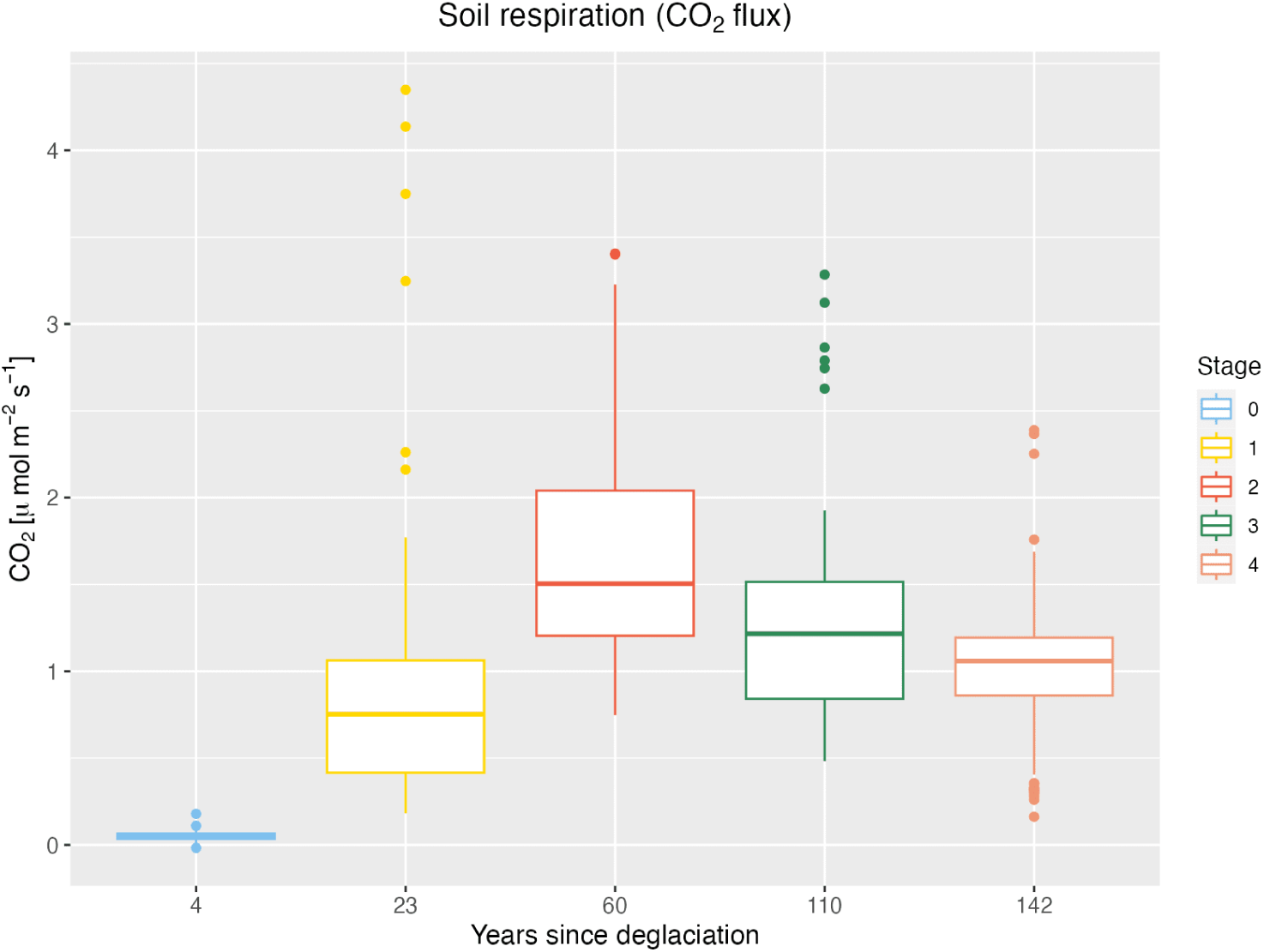
Boxplot of the soil CO_2_ respiration according to glacier retreat. Glacier retreat is represented by years since deglaciation and stages.

**Figure 24:**
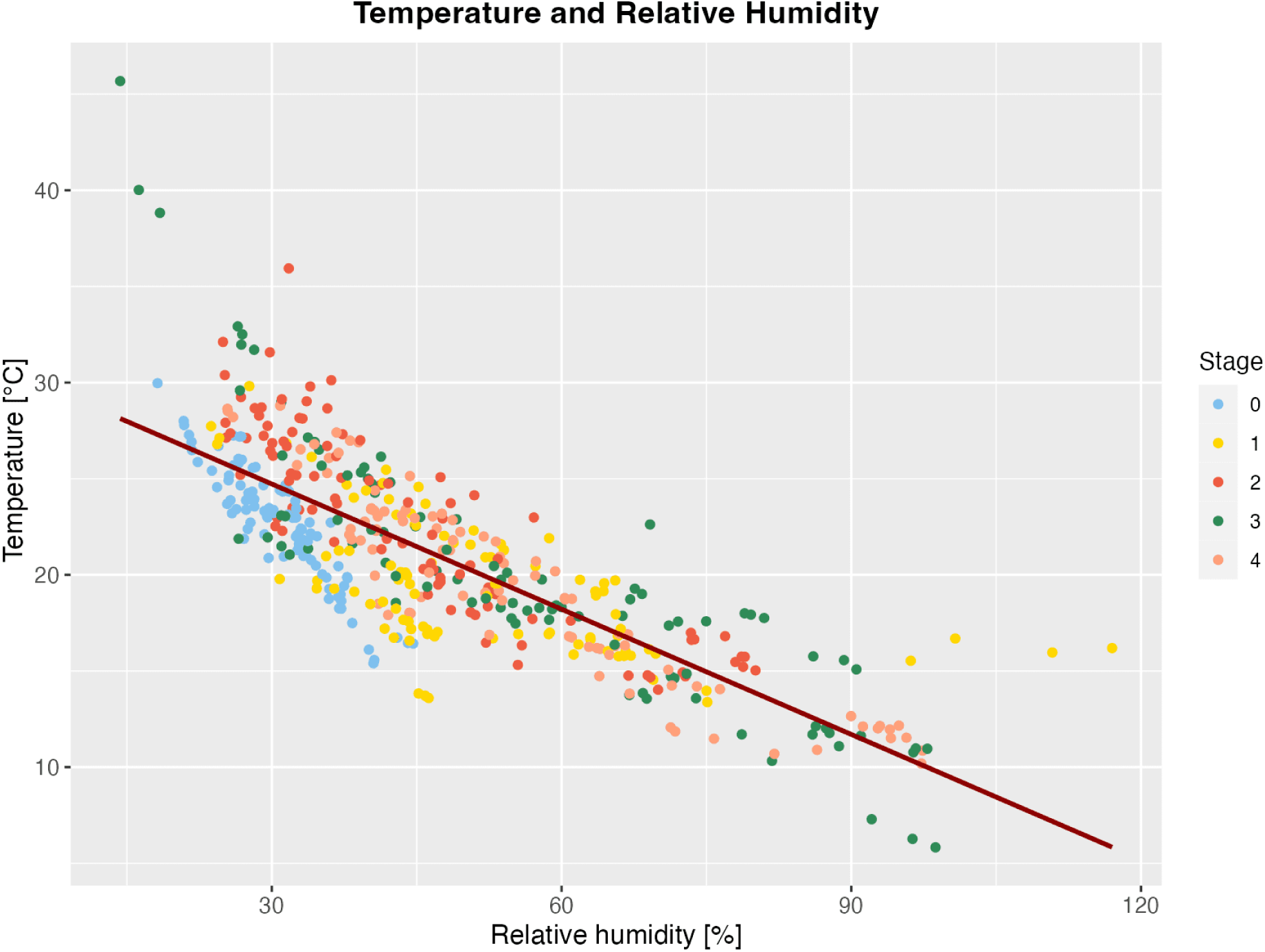
Relationship between temperature and relative humidity. Glacier retreat is represented by stages.

**Figure 25:**
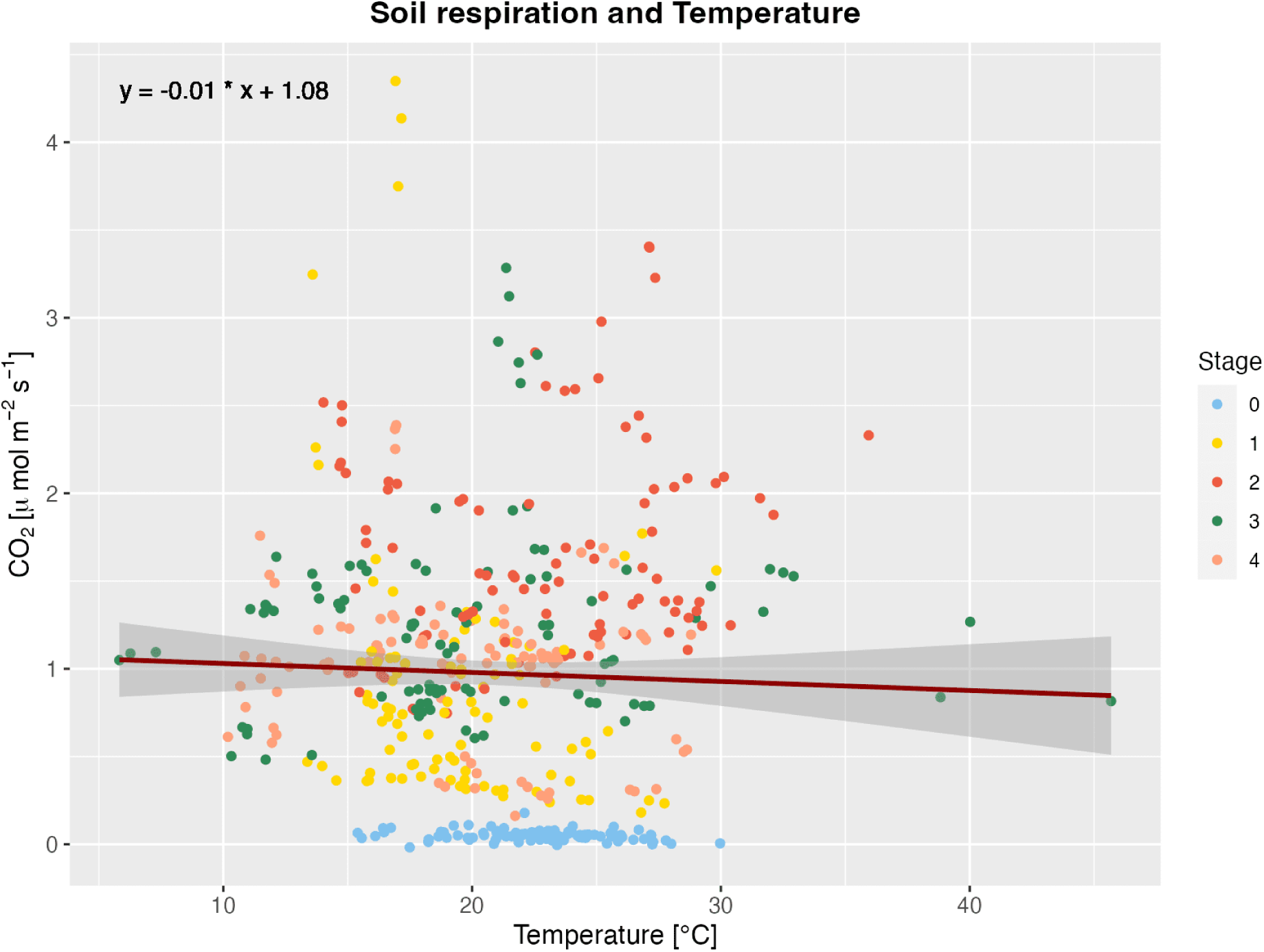
Scatter plot of soil CO_2_ respiration and temperature changes according to glacier retreat. Glacier retreat is represented by years and corresponds to stage 0 (4 years) to stage 4 (142 years).

### Principal Component Analysis

The PCA graph provides a visual representation of the data structure, as illustrated in Figure 26. The first dimension (Dim1) accounts for 49.3% of the variance, and the second dimension (Dim2) explains an additional 19.3% of the variance. In detail, Dim1 mainly explains variables related to vegetation characteristics, including plant diversity, plant richness, and the distribution of plants across different groups. Additionally, Dim1 also explain soil properties such as available nutrients, soil organic matter content, as well as nitrogen and carbon content. On the other hand, the second dimension (Dim2) is more closely associated with variables linked to the geological aspects of the site. This includes measurements of total elements present and values related to soil respiration, providing insights into the site’s geological composition and ecosystem dynamics.

**Figure 26:**
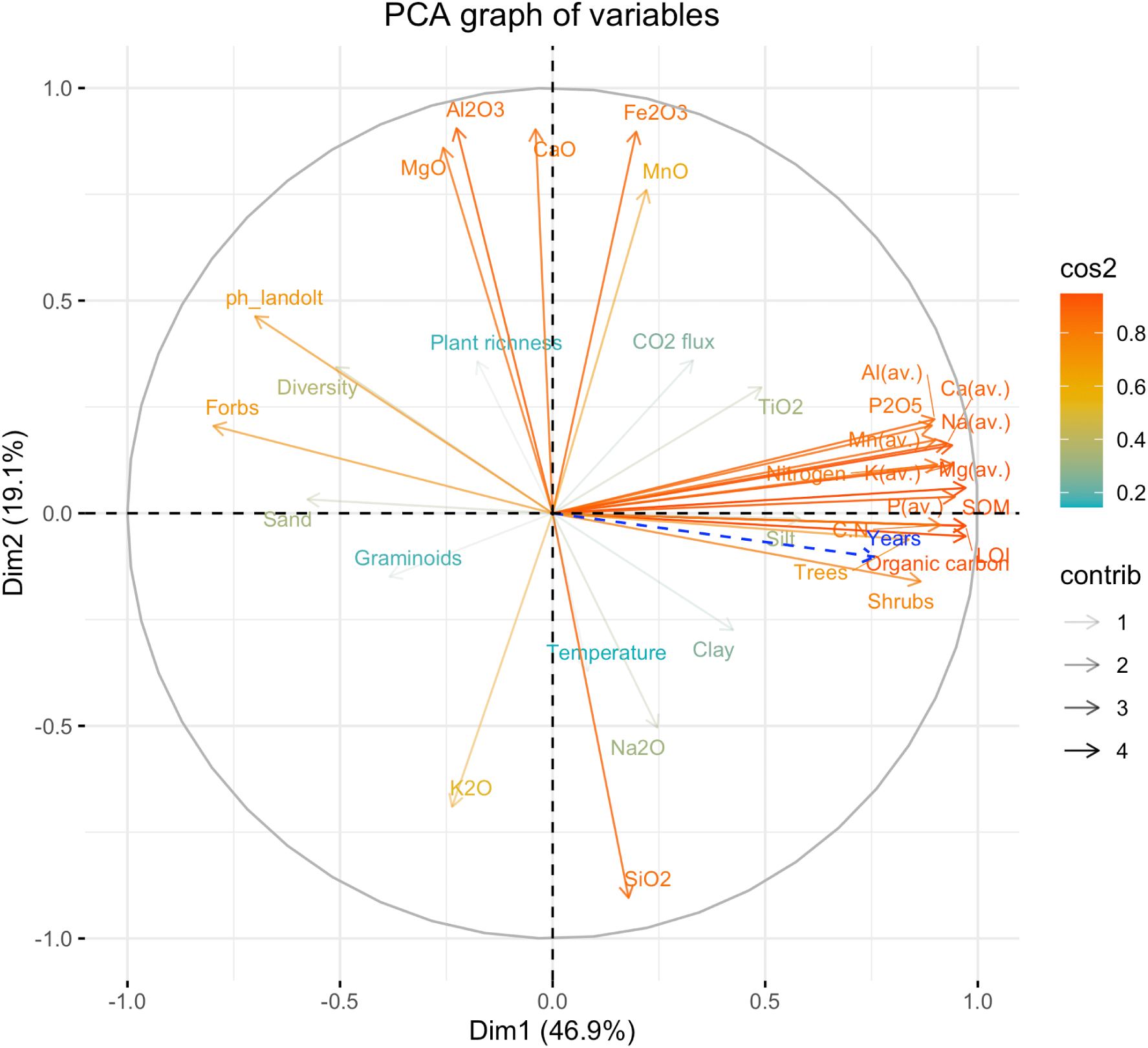
Principal Component Analysis (PCA) graph of variables with squared cosine (cos2) and contribution (contrib) of each variable. Cos2 shows the importance of a component for a given variable. A cos2 value close to 1 shows a better representation of the variable by the principal component. Contrib represents the contribution of each variable to a component. The higher the contribution, symbolized by a darker arrow, the more the observation contributes to the component

Regarding the correlations (Figure 27) among the variables, soil organic carbon exhibits a positive correlation with various available elements (denoted as "av."), including Ca, Mg, Al, K, Na, P, and Mn. Additionally, soil organic carbon is positively correlated with total nitrogen and the C/N ratio. Correlation rates are over 0.75 for soil organic carbon and total nitrogen with all the available elements. On the contrary, pH shows a negative correlation with soil organic matter, soil organic carbon, and available nutrients, indicating a general trend of negative association between pH and nutrient availability in the soil.

**Figure 27:**
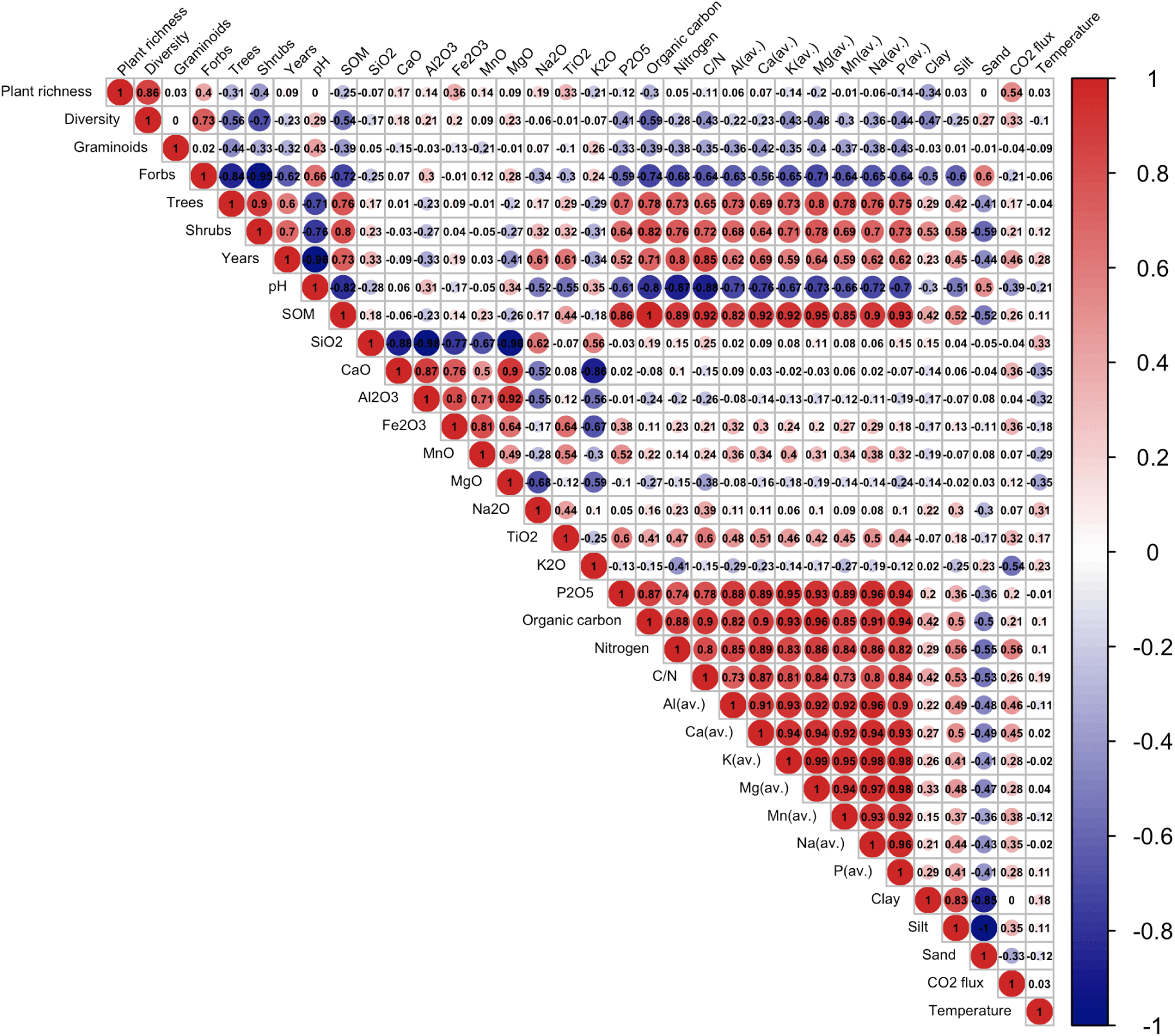
Correlation matrix for all dependent and independent variables. Colors indicate positive (red) or negative (blue) correlation. The areas of the circles indicate the absolute value of the corresponding correlation coefficients.

### 3.4 Structural Equation Modeling (SEM)

Finally, a multivariate statistical analysis model provides an overview of the complex relationships between glacier retreat, vegetation development, soil properties and soil respiration (soil CO_2_ flux) (Figure 28).

**Figure 28:**
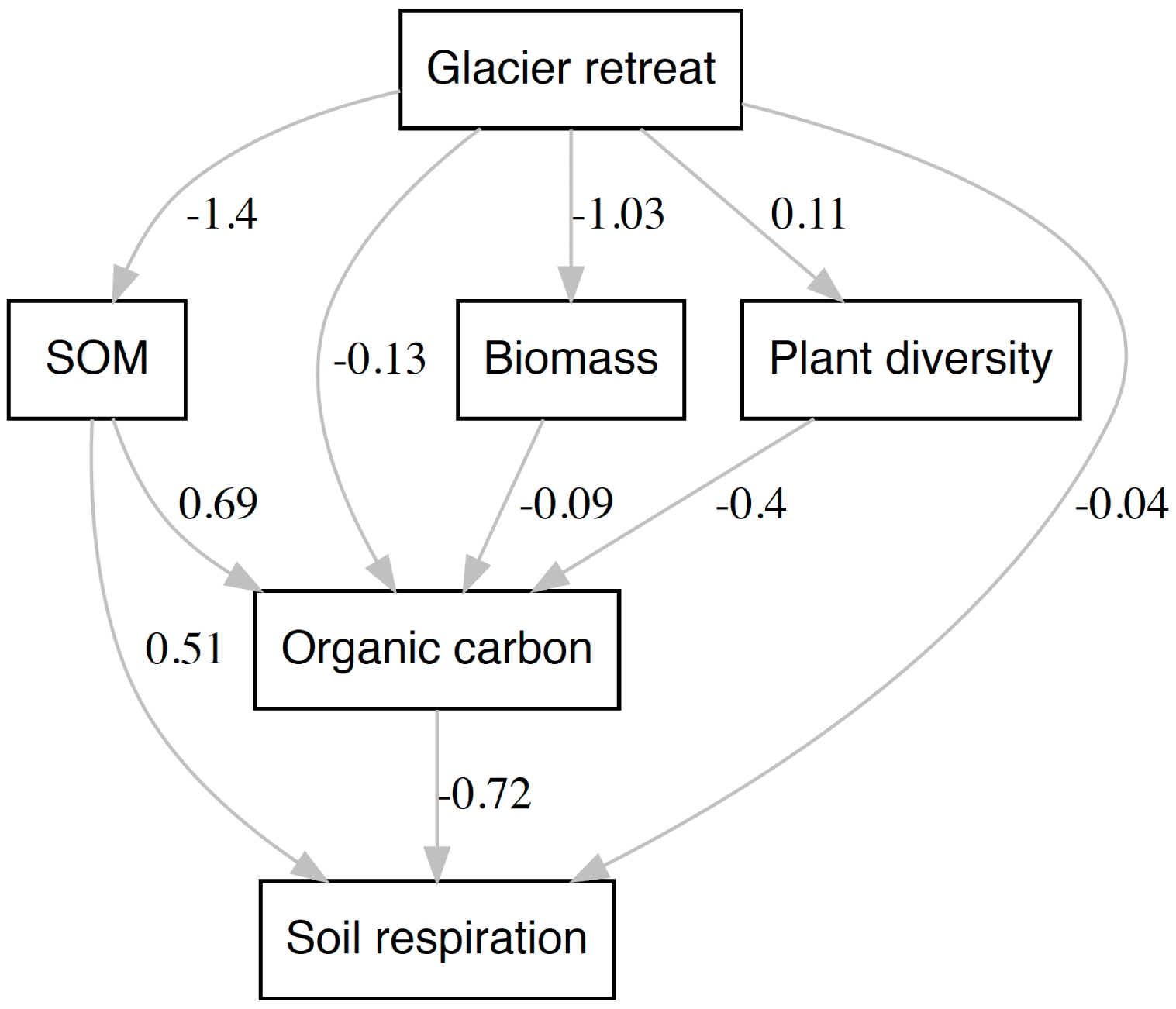
Path analysis showing path coefficients of glacier retreat contributing to changes in soil and vegetation properties and the relationship between the two. Soil properties are characterized by soil respiration (soil CO_2_ flux), soil organic carbon (organic carbon), SOM; vegetation by plant diversity and plant biomass, and glacier retreat by years since deglaciation.

**Figure 29:**
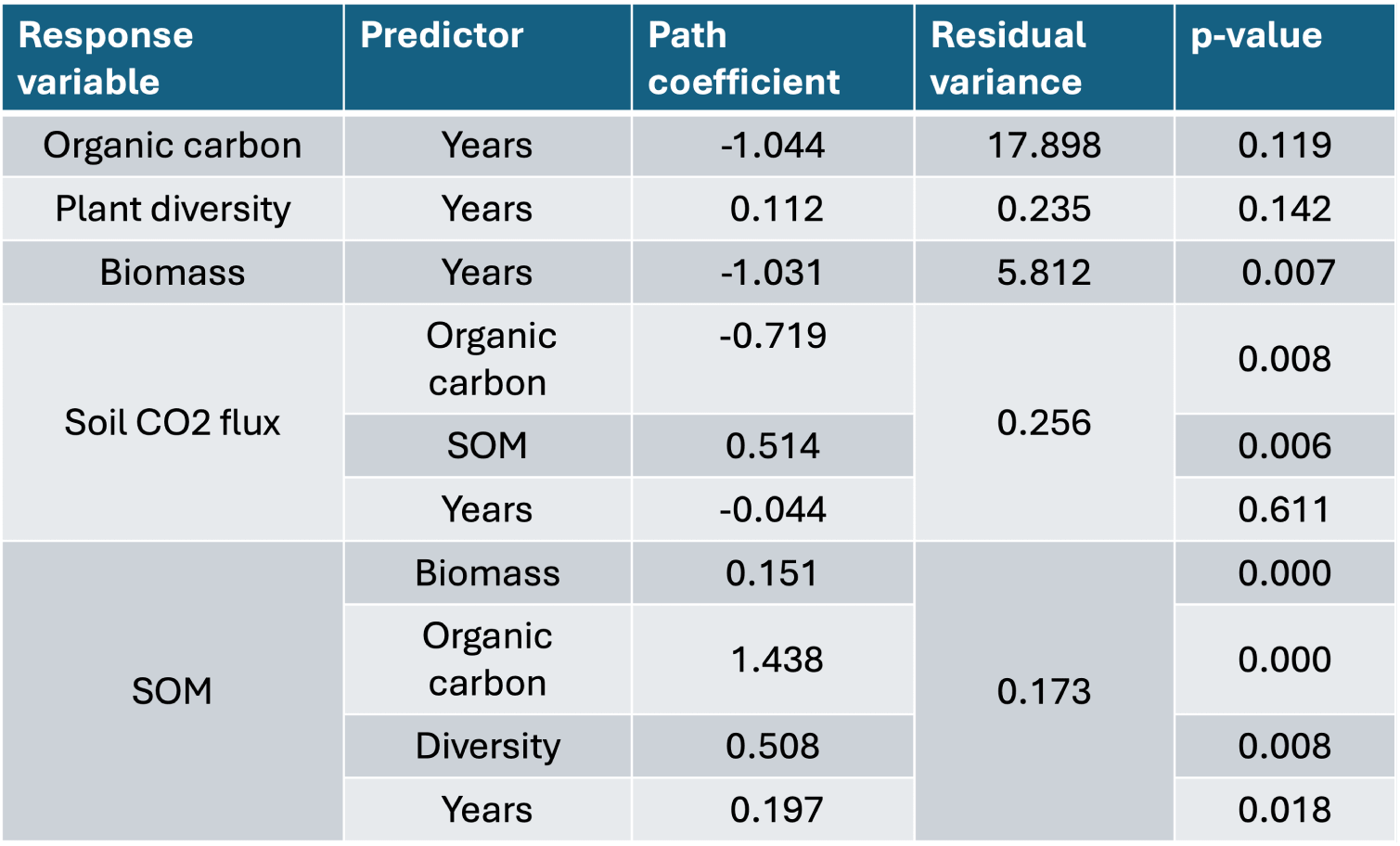
Regression coefficients for path analysis. Path coefficient are regression coefficients that measure relationship between two variables, indicating how many standard deviations of one variable are associated with one standard deviation of the other variable.

**Figure 30:**
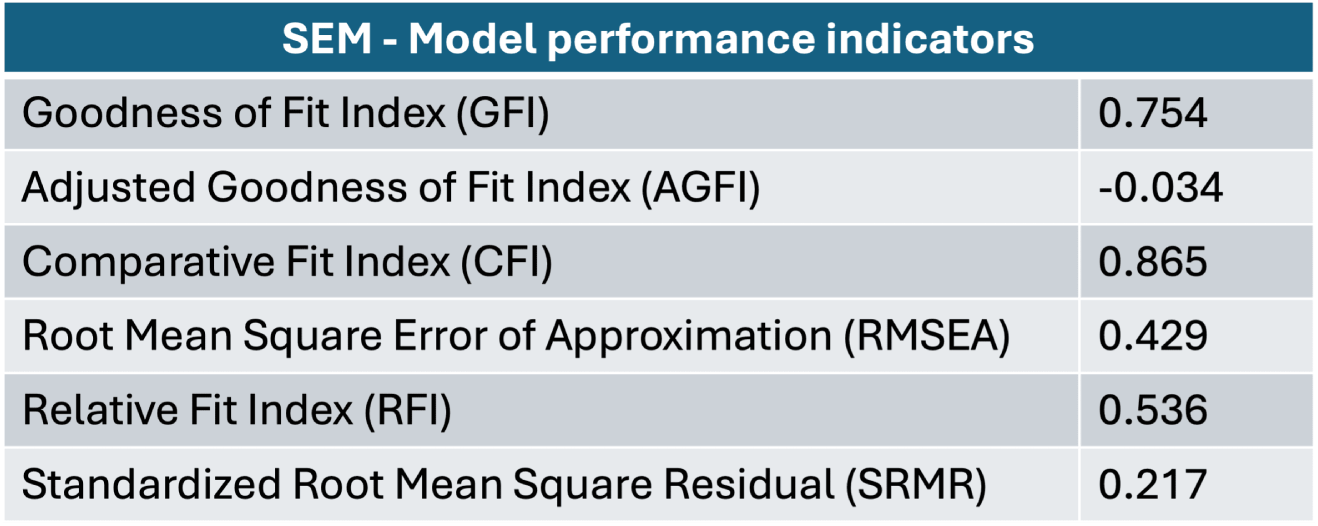
SEM model performance indicators

These models provide a good fit to the data (Table 30). From the model, years since deglaciation is a good predictor of the distribution of soil organic carbon, SOM, plant diversity, biomass and soil respiration (Table 29)

## 4. Discussion

### 4.1 Vegetation and plant diversity

As pioneer plant species begin to colonize barren landscapes, they initiate the process of primary succession. Pioneer species have high adaptability to resist and establish on shallow, nutrient-poor soil composed of rocky debris (Eichel et al., 2013, Bernasconi et al., 2011). At the beginning of the primary succession, abiotic factors are the most important influence on the vegetation dynamics (Ficetola et al., 2021).

The pioneer stage of the succession is colonized by plant communities mostly belonging to the *Epilobietum fleischeri* and the *Oxyrietum digynae* (Burga et al., 2010, D’Amico et al., 2014), and with species belonging to the *Saxifraga* genus (Kabala & Zapart, 2012), which begin the colonization of the raw till, where species have to face establishment on gravel and sandy soil. Pioneer species are highly adapted to specific conditions, developing strategies to cope with substrate and instability disturbances, lack of water and nutrients, and intense sunlight (Eichel et al., 2013, Eichel, 2019). We find that the colonization of plants and shrubs is progressing at a much faster pace than expected. A small shrub, a willow, has already been found at the glacier’s forefront, just 4 years after the glacier disappeared.

Free from ice since 1989, stage 1 represents the early colonization stage, characterized by an increase in vegetation cover (Kabala & Zapart, 2012), plant richness, and diversity (D’Amico et al., 2014). The early stage is enriched in a few species belonging to the Fabaceae genus, which through mutualistic interactions with bacteria, fix nitrogen in soil. Thus, those plants play an important role, in favoring and promoting environmental conditions for late colonizers species, enriching the soil in nutrients and organic matter (Ficetola et al., 2021, Chapin et al., 2002). Here, in addition to nitrogen-fixators plants few young woody pioneer species (i.e., *Larix decidua*, *Salix sp.*, and *Betula pendula*), are found, which usually establish 15 years after glacier retreat (Burga et al., 2010). As late colonizers establish, interactions within plant communities transition from facilitation to competition, favoring generalist species over pioneer specialists (Ficetola et al., 2021).

The intermediate stages, stages 2 and 3, are characterized by the highest level of plant diversity and richness. The coexistence of pioneer species and new species results in maximum species richness and Shannon diversity index. The presence of shrubs and trees increases, and graminacee and forbs remain abundant. Moreover, increasing evidence suggests that mycorrhizal fungi drive plant population biology and community ecology by affecting dispersal and establishment and regulating plant coexistence (Tedersoo et al., 2020).

However, 140 years after deglaciation, in the late successional stage, plant diversity and richness decline. This is due to the dominance of a few species which causes species loss (Hillebrand et al., 2008). The climax forest is composed of the dominant *Larix decidua* and *Picea abies* (Burga et al., 2010). The understory vegetation consists of a few graminacee and forbs, with high coverage of a tick layer of shrubs (i.e., *Rhododendron ferrugineum*, and *Vaccinium sp.*). Additionally, high shrub cover not only causes a decrease in plant species richness but also leads to higher homogenization of the species composition (Pornaro et al., 2017).

The observation of low plant richness and diversity aligns with the findings of Losapio et al. (Losapio et al., 2021), who similarly demonstrate that after a period of saturation and stasis (which happens in the intermediate stage), plant diversity will start declining substantially, losing half of the species across communities, ultimately causing biodiversity loss.

In summary, glacier retreat influences plant community dynamics. First, pioneer colonizers are replaced by more competitive, slow-growing species and tolerant species, suggesting a major role for competition rather than facilitation among plant communities (Ficetola et al., 2021, Mavris et al., 2010). Thus, glacier retreat creates environmental conditions that disadvantage pioneer species, creating an environment that promotes competition and success of late colonizers. Overall, species distribution is explained not only by glacier retreat, but also by species association, and soil properties such as organic matter, gravel, and pH (Losapio et al., 2021).

### 4.2 Soil development

The factors that contribute to soil development are broad, from the nature of the parent material, to topography, climate, and colonization of organisms and time Burga et al., 2010, Egli et al., 2006, Ficetola et al., 2021, Khedim et al., 2021). The present work illustrates how soil properties change along a succession of glacier retreat and how affect soil development.

Stage 0, which corresponds to ground free from ice since 2017, represents the onset of soil development, undergoing the initial processes of physico-chemical and biological weathering. Physico-chemical weathering involves the breakdown of rocks and soils and the chemical reaction of water and atmospheric gases (Sverdrup, 2009), whereas the responsible for most of the biological weathering and transformation are soil microbes (Schulz et al., 2013). Just a few pioneer plants establish themselves at stage 0, which contribute to the initial concentration of organic carbon, total nitrogen, and organic matter, therefore the low values of these parameters (respectively 0.034%, 0.014% 0.11%) are not surprising (Burga et al., 2010, Dümig et al., 2011, D’Amico et al., 2014). The pioneer stage is also characterized by very low availability of nutrients (K, P, Ca, Mg, and Mn), with concentration approximately close to 0, and the highest proportion of elements belonging to silicon (SiO_2_) and aluminium (Al_2_O_3_). The presence of these nutrients is assumed to derive from metamorphic rocks such as orthogneiss and metagabbro. These rocks are rich in ferromagnesian minerals like feldspar or quartz, and pyroxene or plagioclase, respectively (Lambiel et al., 2016). The weathering of these rocks releases minerals that contain elements such as iron, silicon, calcium, sodium, potassium, magnesium, aluminum, manganese, and phosphorus over time (Chapin et al., 2002). These minerals can then release ions that are available to plants as nutrients or remain in a mineral form that cannot be absorbed by roots. Regarding soil texture the pioneer stage is classified as sandy granulometry. This stage, being the most recent ice-free ground, consists of glaciofluvial gravels and sands undergoing the initial process of weathering. Finally, due to the absence of an organic component acidifying the substrate, the pH value is 7.1, corresponding to neutral values.

Located in the 2017-1989 moraines, stage 1 shows the initial formation of vegetation communities and soil development (Egli et al., 2006). This stage represents the early colonization stage, still characterized by low primary production, but with increased plant cover and plant richness. Stage 1 is also characterized by initial soil development. For instance, soil organic carbon, total nitrogen content, and soil organic matter increase, given by the initial input from plant residues (Dümig et al., 2011, Kabala & Zapart, 2012, Burga et al., 2010). Those parameters represent a key element of ecosystem development in the early stages of succession. Over the course of 30 years, due to the continual rock weathering, the concentration of nutrients in the soil slowly increases. The distribution of the elements is associated with weathering processes primarily controlled by climate and vegetation (Kabala & Zapart, 2012). The process of weathering is accelerated by the release of organic acids and the associated pH decrease, due to vegetation establishment (Mavris et al., 2010).

Further away from the glacier front, with increased terrain age, stages 2 and 3 represent the intermediate stages, respectively free from ice for 60 and 110 years. Both stages exhibit increase in plant cover and productivity, leading to a higher transfer of organic matter to the soil (Burga et al., 2010). Organic carbon and total nitrogen content in stages 2 and 3 are similar, reflecting similar vegetation indices in these stages. Indeed, as previously discussed, stages 2 and 3 show the highest plant richness and diversity. The higher accumulation of dead organic matter in soil explains the trend of increased soil organic carbon and total nitrogen. Moreover, SOM accumulation plays a crucial role in soil development as it drives the initial formation of soil structure over time after glacial retreat, forming microaggregates with fine mineral surfaces (Schweizer et al., 2018, D’Amico et al., 2014). Plants, through their root systems and organic matter deposition, contribute to the accumulation of soil organic matter. With an increased input of organic matter, decomposition rates rise, releasing weak acids such as humic acid and fulvic acid (Pettit, 2004), thus lowering soil pH. The decomposition of organic matter not only releases organic acids, but also releases nutrients into the soil, such as nitrogen, phosphorus, potassium, and other essential elements for plant growth (Chapin et al., 2002). As the plant community diversifies and more species establish themselves, the nutrient cycling becomes more efficient (Miransari, 2013). This results in an overall increase in nutrient availability in the soil, creating a positive feedback loop which in turn supports the growth of more diverse and complex plant communities (Chapin et al., 2002, Ficetola et al., 2021).

Finally, the late successional stage, stage 4, is composed of a climax forest, dominated by *Larix decidua* and *Picea abies*, with an understory vegetation dominated by *Rhododendron ferrugineum* and different species of *Vaccinium*. Stage 4 is the one that most differs from the other stages. After 140 years since deglaciation, the soil is developed (Burga et al., 2010, Khedim et al., 2021, D’Amico et al., 2014), due to the continuous biological and psycho-chemical weathering, along with the contributions of organic matter from plants (Bernasconi et al., 2011). Hence, stage 4 shows the highest content of organic carbon, total nitrogen, and nutrients, because of the high vegetation input (Egli et al., 2012, Khedim et al., 2021, Lange et al., 2015). The effect of vegetation on pedogenesis could be enhanced by mycorrhizal fungi associated with different plant species (Burga et al., 2010). In fact, under coniferous or ericaceous species, ectomycorrhizal and ericoid (associated with *Ericaceae*) fungi increase the weathering rate in surface mineral horizons. Moreover, the decomposition of coniferous plant material, such as needles and cones, occurs at a slow rate due to their high lignin content, leading to the build-up of organic matter in soil. As a consequence the superficial soil is enriched in an acidifying litter (D’Amico et al., 2014, Mavris et al., 2010), hence the soil pH value goes toward low value. Soil pH is important as it affects the availability of nutrient elements for plant uptake. With the soil pH falling below pH 6.0, the availability of N, P and K, becomes increasingly restricted. Additionally, soil organic matter stabilization further enhances soil development (Khedim et al., 2021, Schweizer et al., 2018).

Regarding the proportion of elements found in soil, the trend over the ecological succession remains mostly stable. This is due to the time required for the nutrients to be released by the mineral aggregates in soil, which requires more than 150 years. Calcium (CaO) and magnesium (MgO) oxides are the only elements that halve the concentration, due to the dissolution process of carbonate rocks. For the other elements, the difference in concentration is given by input from different sources and different mineralogic substrates (Chapin et al., 2002).

To summarize, going downstream from the glacier’s front in 140 years soil undergoes processes of weathering which lead to soil development. Development proceeds through stages characterized by increase in organic matter content, organic carbon, nitrogen, and nutrients. Moving further away, the slow decomposition of coniferous plant material contributes to an accumulation of organic matter, enriching the soil with acidifying litter. This, coupled with the stabilization of soil organic matter, promotes further soil development, evidenced by increasing SOM content and potentially higher pH levels as vegetation establishes and enriches the soil.

### 4.3 Soil respiration

Soil respiration is a crucial component of the carbon cycle, as it represents one of the primary ways in which carbon is returned from living organisms and plant material back to the atmosphere (Chapin et al., 2002). Soil respiration is carried out by heterotrophic soil organisms (e.g. bacteria, fungi, invertebrates), autotrophs (e.g. plant roots and their associated mycorrhizae), through the degradation of organic matter (Grand et al., 2016) and consequently CO_2_ release.

The soil respiration measurements carried out in the present study have enabled us to characterize the amount and the effect of the CO_2_ flux along the primary succession of glacier retreat. We expected soil respiration to increase with glacier retreat due to the increasing colonization of vegetation and the increase in soil organic matter, microbial biomass, and root biomass. However, our findings show a pattern where the CO_2_ flux increases from stage 0 to 1 to 2, eventually reaching its peak, only to then decrease during stages 3 and 4. This outcome is particularly insightful as it stands in contrast to our initial expectations. For instance, also Wookey et al. (Wookey et al., 2002) observed higher soil CO_2_ effluxes close to the glacier front, with soil respiration rates decreasing with increasing soil age, whereas other studies (Nakatsubo et al., 2005) found soil CO_2_ efflux rates increasing with the time since deglaciation.

Indeed, the measured CO_2_ flux at stages 0 and 1 is very low, which can be explained by the shallow soil depth at the glacier front (Dümig et al., 2011), the non-developed soil (Dacal et al., 2022), and the few plants established with no microbial activity. Afterward, the observed peak in soil respiration at stage 2 corresponds with a simultaneous peak in plant richness. This trend is also supported by the study conducted by Dias et al. (Dias et al., 2010), which found that an increase in soil respiration is closely associated with high plant richness. Indeed, the high number of plant species favors high productivity, which contributes to higher input of organic components in soil and in turn, promotes soil organisms’ activity (Prommer et al., 2020, Metcalfe et al., 2011). Higher soil microorganism activity fosters rates of microbial growth and turnover, thus resulting in higher amounts of microbial biomass. Furthermore, also the type of vegetation affects soil respiration (Bernasconi et al., 2011, Lange et al., 2015, Johnson et al., 2008), shaping the presence of diverse microbial communities in the soil (Eisenhauer et al., 2010, Tedersoo et al., 2020). For instance, the presence of Fabaceae in stages 2 and 3 could affect soil respiration, as those plants have a rapid turnover of organic matter and they favor the presence of nitrogen-fixing bacteria (EL Sabagh et al., 2020, Ficetola et al., 2021). Also the *Salix sp*. species present in these stages are associated with ectomycorrhizae (Bernasconi et al., 2011), which enhance soil microbial activity. Additionally, plant roots have been found to promote soil respiration (Wen et al., 2021), suggesting that a larger root network corresponds to increased respiration. On the other hand, the decrease in soil respiration found in stage 4 can be explained by a potential shift in the microbial composition. Although we did not measure microorganisms’ community composition we can presume that under a coniferous forest, soil microorganism diversity is fairly low and dominated by ectomycorrhizal fungi and ericoid mycorrhizal (Burga et al., 2010). Stage 4 is dominated by *Larix decidua* and *Picea abies*, with the ground covered by *Ericaceae* shrubs such as *Rhododendron ferruginous* and *Vaccinium*. This specific vegetation leads to highly acidic soil conditions, low rates of decomposition of organic matter, and slow turnover of nutrients. The slow decomposition of organic matter by microorganisms impacts soil respiration, thus resulting in lower carbon flux. Furthermore, Stage 4, unlike the other stages, is situated within a closed forest canopy, resulting in shade, lower temperatures, and altered site humidity, likely impacting soil respiration and contributing to its reduction. It is recognized that temperature positively influences soil respiration (Chapin et al., 2002, Grand et al., 2016, Nakatsubo et al., 2005), with higher temperatures leading to increased soil respiration rates. Despite this knowledge, our study did not reveal a significant correlation between these variables, on the contrary, we found a very weak and negative correlation between soil temperature and soil respiration.

Overall, we can say that the increase in soil CO_2_ flux expected with glacier retreat is partially observed. The trend corresponds to the increase in plant colonization associated with an increase in soil microorganism activity and root density. The various stages demonstrate variations in carbon emission, indicating different environmental conditions or processes influencing soil respiration. These findings highlight the intricate interaction between temperature, humidity, and environmental stages in influencing soil CO_2_ fluxes.

### Carbon dynamics

The balance between carbon input (primarily plant-derived matter) and output (primarily soil respiration) in soils is of great importance.

With an estimated of 3,000 Pg of carbon stored globally (Tarnocai et al., 2009) soils are the biggest carbon reservoir. Carbon is stored belowground in the form of soil organic matter. Dead plant material, roots, and other organic residues decompose over time, releasing carbon into the soil where it becomes part of the soil organic carbon pool. Soil respiration is one of the major contributors to carbon dioxide released into the atmosphere, has a CO_2_ efflux estimated at 80 Pg C yr*^−^*^1^ (Raich & Schlesinger, 1992), therefore accounting for a big part of the global carbon emissions. However, with global warming the rate of decomposition of soil organic carbon increases, thereby contributing to the increase in atmospheric CO_2_ and finally, global temperatures (Hopkins et al., 2012). Estimates from a model developed by Nissan et al. (Nissan et al., 2023) show that heterotrophic respiration has been increasing since the 1980s at a rate of about 2% per decade globally, and future projections predict a global increase of about 40% in heterotrophic respiration by the end of the century under the worst-case emission scenario. For instance, if we consider the emission from soil respiration in the present study site, the average CO_2_ flux over the month of measurements and overall the plots is 0.974 *µ* mol m*^−^*^2^ s*^−^*^1^. In a vegetative season of 5 months (average vegetative season in the Alps), the carbon emission could reach the total emissions of approximately 153 Mg C for only the valley itself. While this amount may seem modest in comparison to global emissions, it represents a significant contribution within the context of this specific site. Global warming accelerates decomposition of carbon in forest soils, increasing soil respiration. Increased CO_2_ emission from soils is likely to provide a positive feedback to the greenhouse effect (Raich & Schlesinger, 1992), thereby raising atmospheric temperature and consequently contributing to global warming.

Carbon is stored not only in the soil but also in biomass, representing a critical aspect of the carbon cycle (Chapin et al., 2002). Aboveground biomass, such as trees, shrubs, and other vegetation, serves as a significant reservoir for carbon sequestration. Through the process of photosynthesis, these plants absorb carbon dioxide from the atmosphere, converting it into organic matter that is stored in their leaves, stems, and roots. This biomass acts as a natural carbon sink, playing a crucial role in mitigating the effects of greenhouse gas emissions (Lin et al., 2023).

Open questions remain on how the fragile alpine ecosystems will respond to rising temperatures, higher atmospheric CO_2_ concentrations, shifting seasonal patterns, varying water balances, and nutrient availability. This is particularly significant for ecosystems under threat due to glacier retreat. These environmental changes pose significant challenges to the alpine biodiversity, potentially impacting flora and fauna distribution, diversity, and overall resilience.

### 4.4 Interaction between glacier retreat, vegetation and soil

The path analysis model provides an overview of the impact of Ferpècle and Mont Miné glaciers retreat on vegetation and soil development. This simplified model shows that glacier retreat is a good predictor of soil and vegetation development.

In particular, the negative relationship between soil CO_2_ flux and organic carbon indicates that as the organic carbon content in the soil increases, there is a corresponding decrease in soil respiration, which aligns with our observations of increasing soil organic carbon and the decline of soil respiration with glacier retreat. Additionally, the positive relationship of soil CO_2_ flux with SOM indicates that higher soil organic matter leads to increased soil respiration, as we found in our study as well. Diversity is positively influenced by years since deglaciation, suggesting that over time, there is an increase in plant diversity following glacier retreat. SOM is positively related to both biomass and organic carbon, indicating that as plant biomass and organic carbon in the soil increase, so does the soil organic matter. Additionally, SOM is influenced positively by plant diversity, suggesting that higher plant diversity promotes soil organic matter accumulation. Lastly, years since deglaciation has a positive influence on SOM, implying that as time passes, there is an increase in soil organic matter. Interestingly, the negative impact of years since deglaciation on biomass implies that as time progresses, there is a decrease in plant biomass. However, the biomass data have been transformed in logarithmic for the model to work, which may have introduced some bias. This model also has a limited number of variables, given that few samples were analyzed. To develop a more complex model, more data need to be collected in the future, as the statistics would be more robust and would have facilitated interpretation of the heterogeneity of certain data.

## 5. Conclusion

The retreat of Mont Miné and Ferpècle glaciers has effects on ecosystem development, soil properties, and carbon dynamics. Through the process of primary succession, barren ground is colonized by pioneer plant, initiating a series of changes that eventually lead to the establishment of diverse plant communities. Our study reveals significant insights into how vegetation and soil properties evolve over time since deglaciation, revealing the interactions within these ecosystems.

The pioneer and early stages of succession are characterized by the establishment of specialized plants adapted to harsh conditions. These early colonizers enrich the soil with nutrients and organic matter, promoting the establishment of later colonizer species and increasing plant diversity. Nevertheless, as plant diversity increases, species interactions shift from facilitation to competition. Intense competition results in a few dominant species, where diversity significantly decreases, as seen in stage 4 corresponding to a closed forest. While forbs and graminacee dominate in the early stages, the establishment of shrubs and trees contributes to a decrease in herbaceous species as one moves away from the glacier.

Between the two extremities of our study site, i.e. in 120 years, there is a significant increase in carbon accumulation in both plant aboveground biomass and soil. This is largely due to the establishment and growth of vegetation, along with the input of plant residues, which contribute to the build-up of soil organic matter and higher nutrient levels in the soil. These factors further promote vegetation colonization and the overall development of the soil. Plant growth and soil microorganisms activity affect soil respiration. Soil respiration initially absent at the glacier forefront, increases with plant colonization, reaching a peak in the intermediate stages (2 and 3), and then declining in the late successional stage. The decline in soil respiration in the late successional stage is likely due to the dominance of coniferous species and the resultant slow decomposition rates by soil microorganisms.

With global warming, glacier retreat, and more barren soil, the increase in soil respiration observed in these ecosystems could contribute to a positive feedback loop, further enhancing the greenhouse effect. In conclusion, the retreat of Mont Miné and Ferpècle glaciers has set in motion a dynamic process of ecosystem development, soil formation, and carbon cycling. From the early stages of pioneer colonization to the climax forest communities, these ecosystems are continuously evolving. The findings of this study provide valuable insights into the responses of alpine ecosystems to glacier retreat, highlighting the importance of considering vegetation dynamics, soil properties, and carbon dynamics in understanding the impacts of environmental changes.

### Future perspectives

Further research is needed to fully understand the complex interactions within these ecosystems and their responses to ongoing environmental changes over a longer time scale. Further studies could also investigate the specific mechanisms driving the observed relationships, such as the role of microbial communities in carbon cycling and the effects of changing climate conditions on these processes. As we face a future of rising temperatures and shifting climate patterns, understanding these dynamics will be crucial for predicting how alpine ecosystems will respond to ongoing environmental changes and developing effective conservation and management strategies.

